# Role of polycomb repressive complex 2 in regulation of human transcription factor gene expression

**DOI:** 10.1101/2021.08.04.455052

**Authors:** Jay C. Brown

## Abstract

The study described here was pursued with the idea that information about control of human gene expression might result from comparing the promoter regions of a highly expressed gene population with that of a related population that is weakly expressed. Systematic differences in the two promoter populations would be candidates for a role in causing the distinct expression levels observed. The results reported here were obtained with human genes encoding transcription factors (TF). Among TF genes, those with wide tissue transcription are strongly expressed while those expressed in one or only a few tissues are weakly transcribed in most cases. The promoter regions of the two populations were compared using the results of ChIP-seq experiments. Results demonstrated that the promoter region of broadly expressed TF genes is enriched in binding sites for POLR2A, a component of RNA polymerase II while promoters of tissue targeted genes are enriched in EZH2, a subunit of polycomb repressive complex 2 (PRC2). It was rare to observe promoters with binding sites for both POLR2A and EZH2 or for neither one. The findings are interpreted to indicate that strong expression of broadly expressed TF genes is due to the presence of RNA polymerase II at the promoter while weak expression of tissue targeted promoters results from the presence of PRC2. Finally, transcription factor families were compared in the proportion of broadly expressed and tissue targeted genes. The results demonstrated that most families possess both broadly expressed and tissue targeted members. For instance, this was the case with 16 of 20 TF families. The results are interpreted to indicate that while individual TFs such as EZH2 may be specific for broadly expressed or tissue targeted genes, this is not a property of most TF families. Most have both broadly expressed and tissue targeted members.

## 1. Introduction

Studies on the control of gene expression have benefitted from recent advances in DNA sequencing and from identification of biochemical systems that result in up- or down-regulation of transcription. Whole genome sequences are now available for a wide variety of organisms including humans, and well-characterized elements of regulatory control include promoters, epigenetic signaling, CpG islands, structured chromosome domains, mRNA splicing and many others [1-4]. In view of the variety of regulatory pathways found in higher vertebrates, it is unreasonable to assume that there is an overall regulatory system that applies to all genes. More likely is the idea that distinct regulatory regimes apply to distinct gene populations with the effects of individual regulatory systems integrated to create the overall developmental program observed.

In view of the environment of regulatory studies described above, it is reasonable to suggest that progress in understanding regulatory control might be made by comparing the control elements of two populations of related genes that differ in their level of expression. The idea is that such a comparison might lead to information about the identity of the regulatory features involved, at least in a restricted group of genes.

Here I describe the results of a study employing the strategy described above and focused on the genes encoding human transcription factors (TF). Transcription factors expressed in a wide range of tissues (broadly expressed genes) were found to be highly expressed compared to most tissue targeted TF genes where transcription is weaker. TF expression levels were assessed by the results of RNA-seq experiments while ChIP-seq studies were examined to identify protein binding sites in the gene promoter.

The results are interpreted to support the view that polycomb repressive complex 2 (PRC2) plays a central role in regulating the expression of human TF genes, particularly those of weakly expressed tissue targeted genes. PRC are multi-subunit molecular complexes able to bind the promoter region of a target gene and suppress the gene’s expression [5-6]. First discovered in Drosophila in 1947 [7], PRC functions have been actively studied ever since. PRC or components of PRC are found in all metazoans and in some more primitive species including fungi. Two forms of PRC complex are known, PRC1 and PRC2, and both can introduce covalent modifications into the histone components of chromatin. PRC1 ubiquitinates H2 at lysine119 [8] while PRC2 methylates H3 at lysine27 [9-12]. PRC1 and PRC2 are thought to function together to repress gene expression and/or to report on the expression due to other control factors.

## 2. Materials and methods

### 2.1 Gene databases

#### 2.1.1 All human transcription factor genes (Supplementary Table 1; 1018 genes)

This database was derived from the list of Vaquerizas et al. [13] with a modification involving zinc finger transcription factor genes. ZNF genes were examined individually using GeneCards (https://www.genecards.org) and NCBI Gene (https://www.ncbi.nlm.nih.gov). A ZNF gene was retained in the database only if its role as a transcription factor was supported by an experimental study.

#### 2.1.2 Sample of all human genes (SupplementaryTable 2, 183 genes)

A sample of all human protein coding genes was accumulated to serve as a control for analysis of transcription factor genes. Included genes begin with ACO1 and proceed rightward on chromosome 9 for a total of 183 protein coding genes.

#### 2.1.3 Sample of testis-specific genes (Supplementary Table 3, 204 genes)

This database was curated by Brown [14].

#### 2.1.4 Database of all human transcription factor genes showing predominant structural features present in the TF protein (Supplementary Table 4, 1018 genes)

Structural element annotations were obtained from GeneCards and NCBI Gene.

### 2.2 RNA-seq and ChIP-seq results for transcription factor and control genes

Transcription levels for all genes reported here were obtained from the GTEx Portal of RNA-seq results as reported in the UCSC Genome Browser (version hg38 (https://genome.ucsc.edu/)). A gene was annotated as “broadly expressed” (B in Supplementary Table 1) if it was found to be expressed in 90% or more of the 52 tissues reported by GTEx. Otherwise, the gene was annotated as “tissue targeted” (T). Borderline genes were rare (10 of 1018 genes). They are indicated by B/T or T/B in Supplementary Table 1 depending on the category they most resemble. Transcription levels for broadly expressed genes were the average of the top 5 tissues reported by GTEx. For tissue targeted genes, reported values were those of the tissue specific gene or the average of the top 5 tissues in cases where more than one tissue was in the group.

Transcription factor binding sites in the promoter region of transcription factor genes were derived from ChIP-seq studies reported in the ENCODE project database of cis-Regulatory Elements (3 November 2018 version) by way of the UCSC Genome Browser. Visual scanning of promoter binding sites was done with UCSC Genome Browser. An EZH2 entry was included in Supplementary Table 1 if it was present in 50% or more of the studies reported by ENCODE. Values for the level of EZH2 promoter binding were those reported by the Integrated Genomics Viewer (https://igv.org/app/); the cells and study were GM12878 and ENCFF273NDY, respectively. Only binding sites in the promoter region are reported here; sites in non-promoter regions were ignored.

### 2.3 Transcription factor families and protein structural features

These were derived from GeneCards and from NCBI Gene.

### 2.4 Data handling

Data were manipulated with RStudio and Excel. Graphic rendering was done with SigmaPlot v14.5.

## 3. Results

### 3.1 Broadly expressed and tissue targeted human TF genes

The project was carried out beginning with a database of 1018 human transcription factor genes (Supplementary Table 1). The database was that of Vaquerizas et al. [13] with the modifications described in Materials and Methods. Each gene was classified as either broadly expressed or tissue targeted using information about its tissue expression in the RNA-seq results reported by GTEx Portal via the UCSC Genome Browser (https://genome.ucsc.edu/). Of the 1018 database genes, 589 (58%) were classified as broadly expressed and 419 (41%) as tissue targeted (Supplementary Table 1, B/T column).

### 3.2 Gene expression in broadly expressed and tissue targeted TF databases

It was expected that broadly expressed TF genes would be expressed at a higher level than that of tissue targeted TF genes because of the need for expression of tissue targeted TF’s to be suppressed in some tissues. This expectation was tested by rank ordering each population according to the expression level of each gene. The two populations were then plotted together so that the overall expression level could be compared. The results (Fig. 1) showed that tissue targeted TF genes are expressed at a lower level than broadly expressed ones, particularly among the genes with the lowest transcription levels. Above a rank of ∼180, expression of the two populations was more equal with the tissue targeted population exceeding the broadly expressed genes at the highest expression levels (Fig. 1). It was concluded that comparison of the two populations was appropriate for the studies contemplated here.

**Figure 1.**
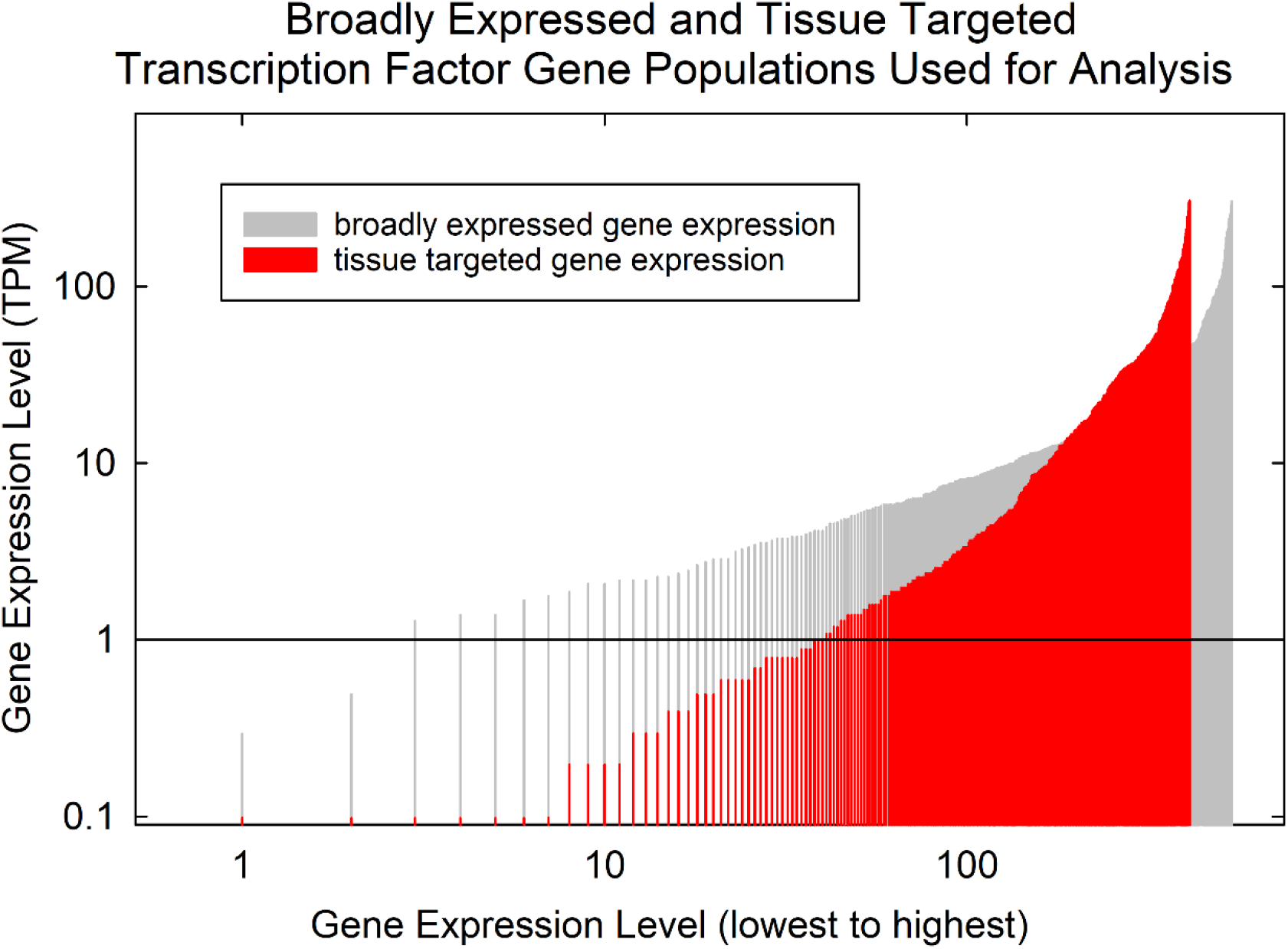
Expression level of broadly expressed compared to tissue targeted human transcription factors. The expression levels of genes in the two populations were ordered by level and compared by plotting on the same graph. Note that the level of tissue targeted TF expression is less compared to broadly expressed genes, particularly in low expression range.

### 3.3 TF binding sites in the promoter region of TF genes

TF binding sites were identified using ChIP-seq results available from ENCODE. At first, identified sites in graphical form were examined visually in the UCSC Genome Browser. Images revealed that binding sites for POLR2A and EZH2 were asymmetrically distributed in broadly expressed compared to tissue targeted TF gene promoters. For instance, among 589 broadly expressed TF genes, 499 (85%) had one or more POLR2A binding sites in the promoter while among 419 tissue targeted TF genes, 246 (59%) had binding sites for EZH2 (Fig. 2a). It was rare to find EZH2 binding sites in broadly expressed gene promoters (7%) or POLR2A in tissue targeted genes (12%; see Fig. 2a). It was also rare to find TF gene promoters that lacked both POLR2A and EZH2. Of the 1018 genes examined, only 54 (5%) lacked binding sites for both (Fig. 2a).

**Figure 2.**
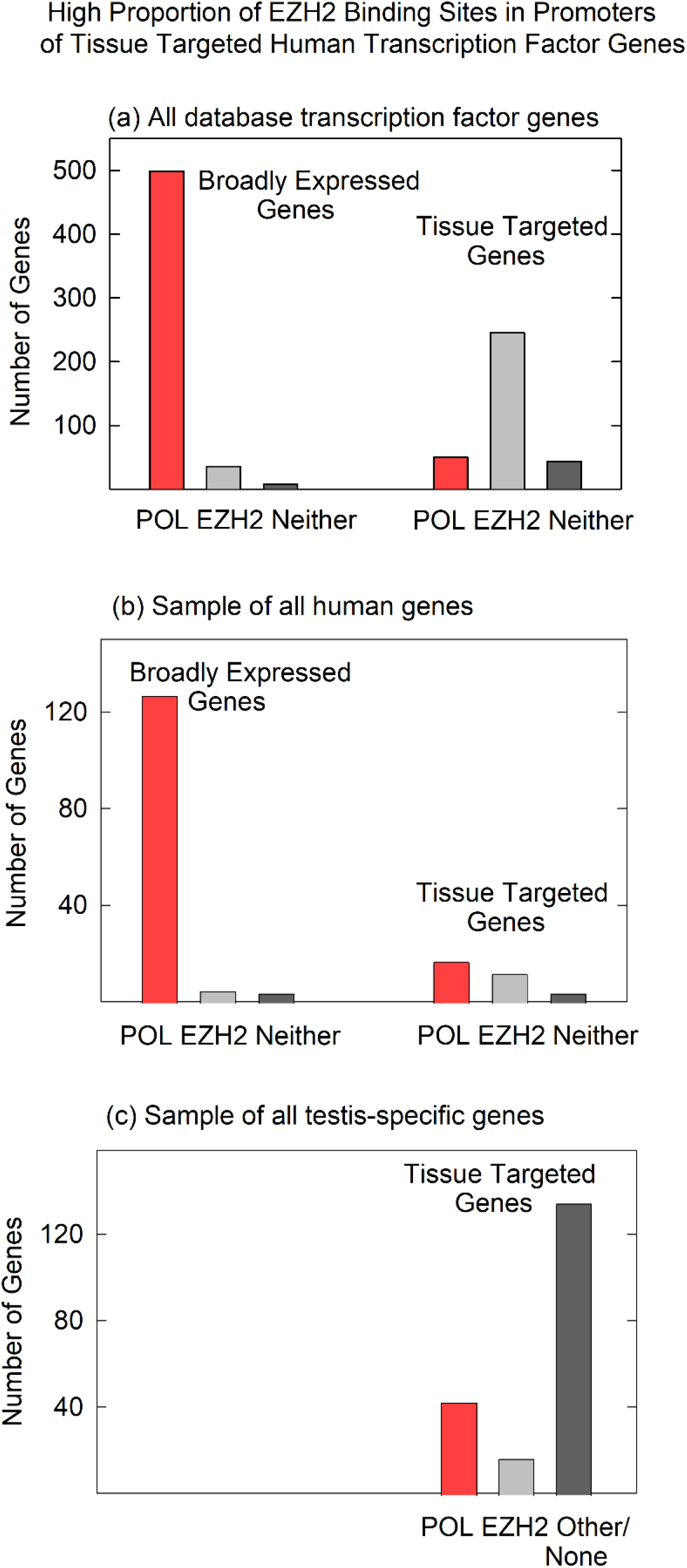
POLR2A and EZH2 binding sites in the promoter regions of broadly expressed and tissue targeted genes. Results are shown for (a) all database human transcription factor genes; (b) an unselected population of human protein coding genes; and (c) a sample of human testis-specific genes. Note that EZH2 binding sites (indicative of PRC2 binding) are prominent in tissue targeted TF genes, but not in broadly expressed TF genes (a), a mixed population of human protein coding genes (b) or in testis-specific human genes (c).

### 3.4 Control: POLR2A and EZH2 binding sites in the promoters of unselected human protein coding genes

As a control, an experiment was performed to test whether the asymmetry observed between POLR2A and EZH2 in the promoter region of TF genes would also be observed in an unselected population of human protein coding genes. Is asymmetry observed only in TF genes, or does it exist more widely? To test the idea, the experiment described above was repeated beginning with an unselected population of all human protein coding genes rather than with TF genes only (Supplementary Table 2). The results showed that among 141 unselected, broadly expressed genes, 127 (91%) had POLR2A in the promoter while 8 (6%) had EZH2, a result comparable to the TF gene population. In contrast, in the tissue targeted population the results were 40% and 29% for POLR2A and EZH2, respectively, a result quite different from the TF genes (Fig. 2b). The results are interpreted to support the view that the POLR2A/EZH2 asymmetry is found in TF genes only and is not found in an unselected gene population.

### 3.5 Control: POLR2A and EZH2 binding sites in testis-specific genes

A further control study was carried out to examine POLR2A and EZH2 binding in the promoters of a population of tissue targeted genes, a population of 204 testis-specific genes (Supplementary Table 3). ChIP-seq results, as described above, were examined in the promoter regions of the testis-specific genes. It was expected that EZH2 binding sites would be observed in a high proportion of genes if testis-specific genes were regulated in the same way found for tissue targeted TF genes. Quite a different outcome was detected (Fig. 2c).

Promoters of testis-specific genes were not found to be enriched in either POLR2A or EZH2. Instead, the promoters had a mixture of transcription factor binding sites in which no predominant species could be identified (see Supplementary Table 3). For instance, of 195 testis-specific genes that yielded a result, 135 (69%) had neither a POLR2A nor an EZH2 site. POLR2A and EZH2 were found in 22% and 9%, respectively (Fig. 2c). The result is interpreted to indicate that while EZH2 binding sites are prevalent in the promoters of tissue targeted TF genes, the same is not found in other populations of tissue targeted genes.

### 3.6 EZH2 represses TF gene expression as a component of the polycomb repressive complex 2

Binding of EZH2 to the promoter regions of tissue targeted TF genes as described above was interpreted to indicate that EZH2 acts to repress TF gene expression as a component of PRC2, a complex known to be involved in gene repression [5-6]. EZH2 is a prominent component of PRC2, and PCR2 introduces repressive modifications into chromatin. To test the idea that EZH2/PCR2 acts to repress TF gene expression, the expression level of tissue targeted TF genes was plotted against the amount of EZH2 reported in the TF gene promoter (Fig. 3). Creation of the plot was complicated by the fact that as EZH2 is repressive, it was difficult to find an appropriate population of tissue targeted TF genes. Most were expressed to only a background level. Fig. 3 shows the results of all 24 of the genes with transcription levels above background. The regression line in the plot indicates that the amount of promoter bound EZH2 is corelated with lower rather than higher gene expression. The results support the conclusion that EZH2 is repressive for its target TF gene population. The results are also compatible with the idea that EZH2 acts as a part of the PRC2 complex.

**Figure 3.**
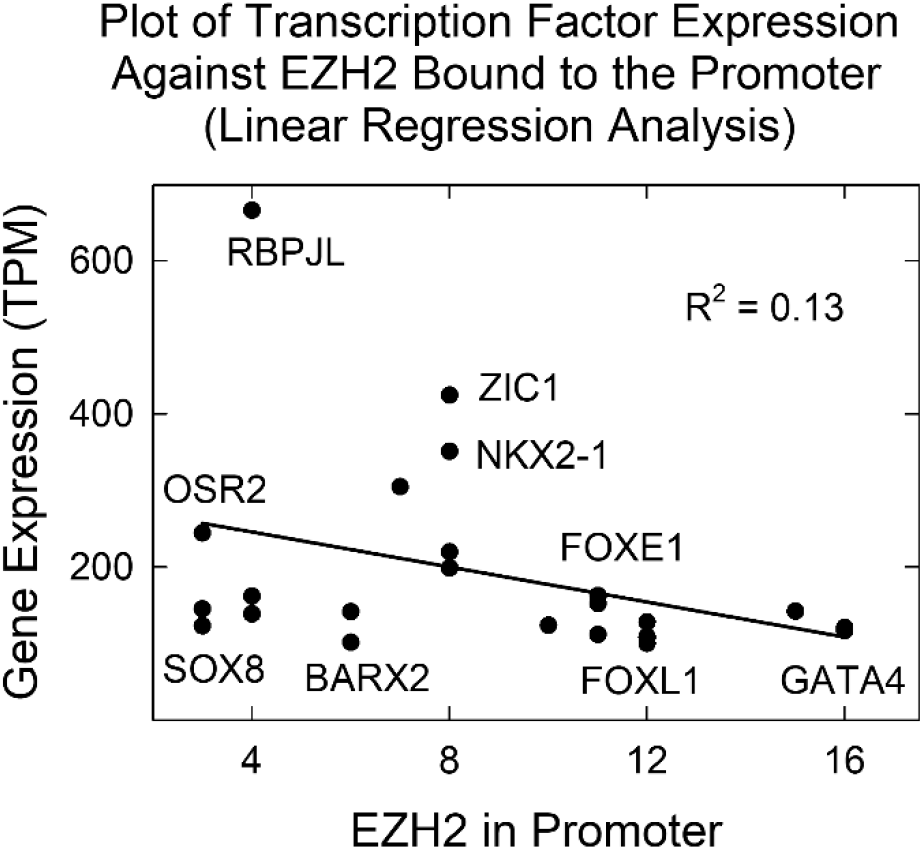
Plot showing transcription factor expression level as a function of EZH2 protein bound to the promoter. Note that increased EZH2 binding correlates with decreased TF gene expression. Data points corresponding to a few individual TFs are noted on the plot.

### 3.7 Preferred levels of EZH2 bound at target promoters

The availability of quantitative values for the level of EZH2/PRC2 in the promoters of tissue targeted TF genes enabled a determination of whether the collection of values is a continuous function or whether there exist preferred levels. The issue was examined beginning with all 246 database TF genes that were judged to be tissue targeted and also to be enriched in EZH2 binding in the promoter (see Supplementary Table 1, EZH2 column). A histogram was plotted relating the amount of EZH2 bound at the promoter and the number of times a similar value was found among the 246 genes (Fig. 4). The plot shows evidence of preferred binding levels in promoter associated EZH2. At least four preferred complexes were identified (at 4,8,12 and 16 EZH2 units), and spacing was found to be ∼4 in the units provided by the ChIP-seq analysis (signal p-value).

**Figure 4.**
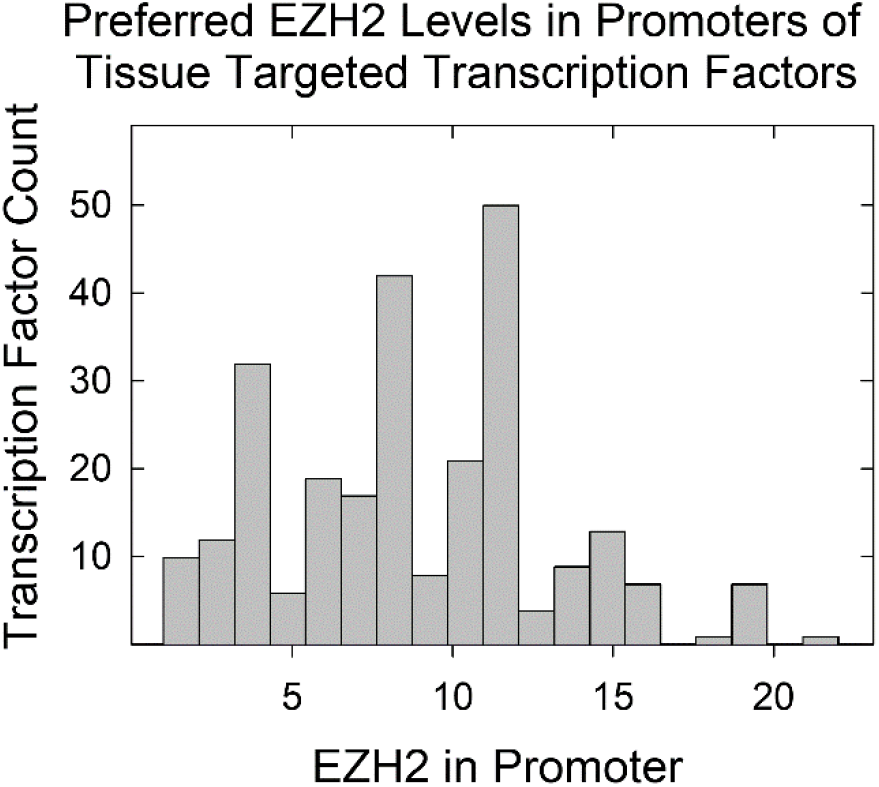
Histogram showing TF gene count as a function of the measured level of EZH2 protein bound at the promoter. Analysis was carried out with all 246 tissue targeted database TFs with EZH2 bound at the promoter. Note that there are preferred levels of EZH2 protein bound.

The above result was not expected. Preferred spacing was not observed when similar histogram analysis was performed with gene expression values from the same dataset. The observation (Fig. 4) is interpreted to indicate that EZH2/PRC2 complexes are added to promoter regions in units containing two or more EZH2 molecules. Preferred spacing arises when more than one multi-EZH2/PRC2 complex attaches to a promoter.

### 3.8 Transcription factor families

The information above indicating that promoter binding sites for EZH2 are located preferentially in tissue targeted TF genes raises the question of whether there may be families of TFs whose members have a selectivity for controlling tissue targeted TF genes only, or broadly expressed TF genes only. Could binding to tissue targeted promoters be a property of a TF family and not only of an individual TF? To explore the above question, the TF family of each TF employed here was recorded and the families were characterized according to the number of broadly expressed and tissue targeted genes they contain. All results are shown in Supplementary Table 4 with a sample of 20 families shown graphically in Fig. 5.

**Figure 5.**
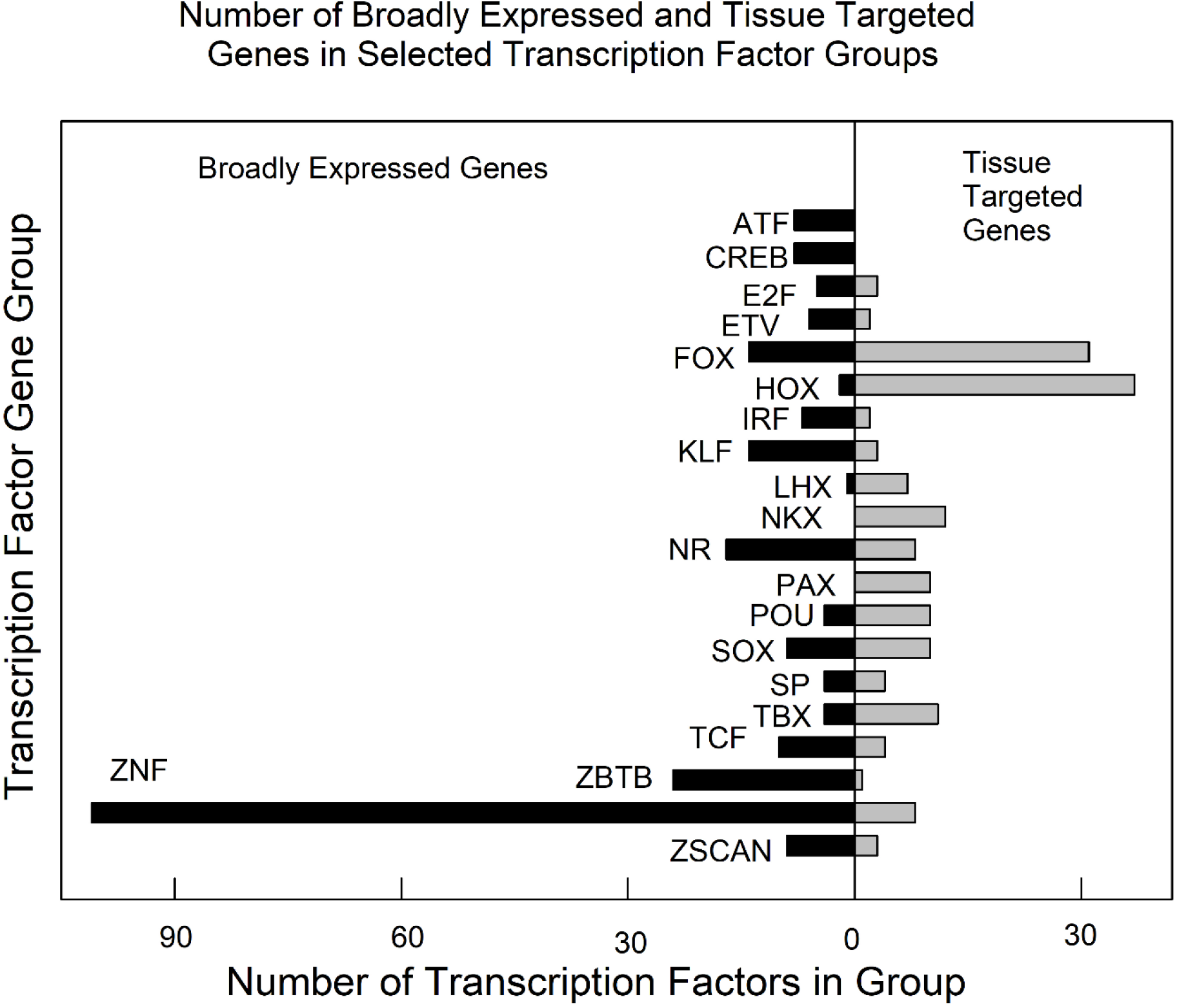
Number of broadly expressed and tissue targeted TFs in selected TF families. See Supplementary Table 4 for all results. Note that most families contain both broadly expressed and tissue targeted TF genes. Families shown: ATF, activating TF; CREB, cAMP response element binding TF; E2F, E2 factors; ETV, FGF-regulated TFs; FOX, forkhead box; HOX, homeodomain; IRF, interferon-regulatory factor; KLF, Krüppel-like TF’s; LHX, LIM homeobox; NKX, NKX homeodomain box; NR, nuclear receptor TF’s; PAX, paired homeobox; POU, Pit-Oct-Unc TF’s; SP, SRY-related HMG box; SP, Sp; TBX, T-box; TCF, TCF/LEF; ZBTB, ZBTB family; ZNF, zinc finger TF’s; ZSCAN, zinc finger and scan domain TF’s.

Of the 20 families, the greatest number (16) contain both broadly expressed and tissue targeted members. The remaining four families have members with only broadly expressed (ATF and CREB) and only tissue targeted (NKX and PAX) genes. Members of the largest family (ZNF; 109 genes) are predominantly broadly expressed genes (101 broadly expressed vs. 8 tissue targeted genes). Forkhead (FOX) and homeobox (HOX) families have the highest number of tissue targeted genes (31 and 37, respectively; see Fig. 5). The results support the conclusion that most human TF families contain both broadly expressed and tissue targeted member genes.

A similar analysis was performed with TFs grouped according to the structural motif found in the TF protein. Eight prominent structural motifs found among TF proteins were identified and the number of broadly expressed and tissue targeted TFs was noted in each one (Fig. 6). The results resembled those observed in family groups. Of the eight structural groups, both broadly expressed, and tissue targeted TF were found to be present in all but one (ZNF-BTB; see Fig. 6). Homeobox were found to have the highest number of tissue targeted TFs and ZNF the largest number of broadly expressed.

**Figure 6.**
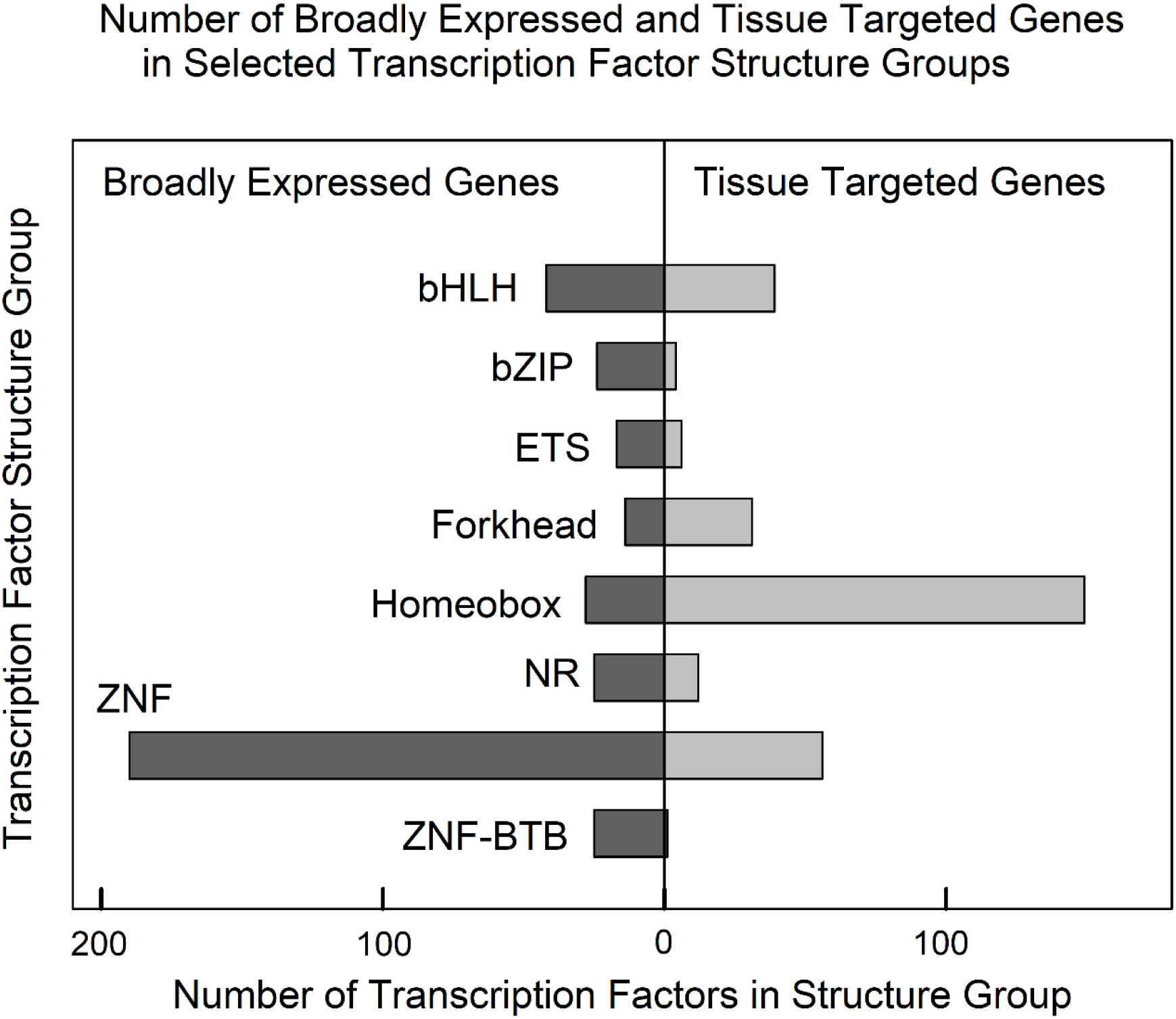
Number of broadly expressed and tissue targeted TFs in selected groups sharing the indicated protein structural feature(s). See Supplementary Table 4 for all results. Note that all structure groups contain both broadly expressed and tissue targeted TF genes. Structural groups shown: bHLH, basic helix-loop-helix; bZIP, basic leucine zipper domain; ETS, erythroblast transformation specific domain; Forkhead, forkhead box; Homeobox, homeobox sequence; NR, multiple structural domains; ZNF, zinc finger domain; ZNF-BTB, zinc finger plus bric-à-brac motifs.

## 4. Discussion

### 4.1 Experimental strategy; comparing weakly and strongly expressing gene populations

The experimental strategy adopted here suggests itself as a pathway to make further progress in understanding the control of gene expression. The idea is to start with two populations of related genes that differ in their level of expression. The results of ChIP-seq experiments are then used to compare TF binding sites in the promoter regions with the goal of identifying sites that differ in the two gene populations. In principle, such a study could begin with any two populations of genes that differ in their level of expression. In practice, however, the best outcomes might be expected with genes resembling each other in features such as function, biochemical pathway, or evolutionary time the genes entered the human genome. In view of the wide range of regulatory systems that could impact a gene, the chance of identifying genes using the same control system might be best in genes that resemble each other in other respects.

### 4.2 Broadly expressed compared to tissue targeted genes

The distinction between broadly expressed and tissue targeted genes is a fundamental one in developmental and evolutionary biology [15-18]. For a gene that might benefit the fitness of the human genome, for instance, if the gene can be broadly expressed then its pathway to incorporation into the genome is simpler than if tissue specific expression is required. In the case of tissue specific expression, the gene needs to enter the genome and be expressed, but further evolutionary steps are required to ensure that the gene is not expressed in off-target tissues. In view of the quite different nature of broadly expressed and tissue targeted genes, it is not surprising that the two gene types have different mechanisms to control their expression as described here. Tissue targeted TF genes were found to have a mechanism, involving PRC2, that is less abundant in broadly expressed TF genes. In fact, it is reasonable to expect that there are other yet to be discovered mechanisms that distinguish gene expression in broadly expressed compared to tissue targeted genes.

### 4.3 High abundance of EZH2/PRC2 in the promoters of tissue targeted TF

An unexpected outcome of the study described here was the high proportion of EZH2/PRC2 found in the promoters of tissue targeted TF. The presence of repressive signals in the promoters was expected [19], but in view of the many types of repressive signals available in the human genome, it was expected that a greater variety would be observed. Instead, of 419 tissue targeted TF examined here, 246 (59%) were found to have EZH2/PRC2 binding sites in the promoter. The high proportion suggests repression by PCR2 is particularly well suited for repression of tissue targeted TF.

Clues about the close link between PCR2 and tissue targeted TF might arise from considering X-chromosome inactivation, a prominent focus of PRC function. PRC2 is well known for its role in suppressing all gene expression from one X chromosome in human females. Targeted to one of the two X chromosomes by an untranslated RNA (Xist), PRC complexes bind along the entire chromosome to silence the genes permanently, creating the Barr body [20-22]. Permanent repression is not observed, however, in all X inactivation situations. Temporary, reversible inactivation has been observed in mouse and in mouse ES cells [23-24]. The case for reversible inactivation therefore needs to be considered open in the case of human TF cells.

PRC2 deposition to create the Barr body differs in another significant way from deposition at promoter regions. Whereas PRC complexes coat an entire X chromosome to form a Barr body, deposition sites are widely distributed in the human genome to suppress tissue targeted TF gene expression [19]. Most tissue targeted TF genes are expected to be separate sites of PRC2 deposition. This situation creates a more formidable problem for targeting complexes at TF genes than it does with an entire X chromosome. Understanding of PCR2 targeting to non-X chromosome sites is further complicated by the ability of polycomb complexes to bind not only to chromatin, but also to RNA molecules as they are being synthesizing [25].

### 4.4 Families of human TF

Evidence that a TF can act selectively on a tissue targeted gene (e.g. EZH2) or a broadly expressed one (e.g. POLR2A) as described here raises the possibility that such selectivity might apply to TF groups as well as to individual TFs. Could all members of a TF family, for instance, be specialized to target only broadly expressed genes? Studies to address this issue generally supported the view that most TF groups contain members able to recognize broadly expressed genes and others that recognize tissue targeted ones. Little evidence was observed for broadly expressed/tissue targeted specificity among either TF families or among TFs with the same protein structural motifs (Figs 5 and 6; Supplementary Table 4). Despite the lack of overall specificity observed, some examples of selectivity were noted. For instance, zinc finger TFs exhibited a strong preference for broadly expressed TF genes while homeodomain TFs showed a preference for tissue targeted genes (Fig. 6).

I suggest that future studies of gene regulatory control might benefit from a focus on TF groups such as NKX and PAX in which all members have tissue targeted expression. It is expected that the promoter regions of these genes will have binding sites for both activating elements able to activate genes in the targeted tissue and repressive elements for non-targeted tissues. Examination of the promoters in such genes might facilitate the identification of the features that influence gene expression.

## Data Availability

All data employed in this study are contained in Supplementary Tables 1-4 from which it can be freely downloaded.

## Acknowledgements

I gratefully acknowledge Karsten Siller for advice on computational aspects of this project. This manuscript was submitted as a pre-print in the link https://biorxiv.org/content/10.1101/2021.08.04.455052v1 [26].

