## Supplementary Table1 for "Role of polycomb repressive complex 2 in regulation of human transcription factor gene expression"

Supplementary Table 1: All database transcription factor genes with information regarding broadly vs. targeted expression, EZH2 vs POLR2A in the promoter; gene expression level and EZH2 binding in the promoter (1018 genes)

| Chr | Gene | B/T | POLR2A/EZH2 | Gene Expression | EZH2 Binding |
|---|---|---|---|---|---|
| 20 | ADNP | B | POLR2A | 44.0 | |
| 18 | ADNP2 | B | POLR2A | 13.3 | |
| 12 | AEBP2 | B | POLR2A | 21.7 | |
| 2 | AFF3 | T | EZH2 | 8.8 | 8 |
| 5 | AFF4 | B | POLR2A | 40.2 | |
| 7 | AHR | B | POLR2A | 49.6 | |
| 21 | AIRE | T | EZH2 | 4.0 | 2 |
| 9 | AKNA | B | POLR2A | 43.8 | |
| 12 | ALX1 | T | EZH2 + POLR2A | 2.6 | |
| 1 | ALX3 | B | EZH2 > POLR2A | 11.7 | |
| 11 | ALX4 | T | EZH2 | 1.1 | 15 |
| 23 | ARX | T | POLR2A + EZH2 | 6.1 | 8 |
| 3 | ARGFX | T | neither | 0.5 | |
| 19 | ARID3A | B | POLR2A | 6.2 | |
| 10 | ARID5B | B | POLR2A | 40.8 | |
| 1 | ARNT | B | POLR2A | 40.2 | |
| 15 | ARNT2 | B | EZH2 + POLR2A | 15.5 | |
| 11 | ARNTL | B | POLR2A > EZH2 | 24.2 | |
| 12 | ARNTL2 | B | POLR2A | 4.1 | |
| 23 | ARX | T | EZH2 | 6.1 | |
| 11 | ASCL2 | T | EZH2 | 3.1 | 6 |
| 11 | ASCL3 | T | neither | 8.7 | |
| 12 | ASCL4 | T | EZH2 | 0.1 | 8 |
| 1 | ASH1L | B | POLR2A | 23.7 | |
| 12 | ATF1 | B | POLR2A | 21.3 | |
| 2 | ATF2 | B | POLR2A | 32.4 | |
| 1 | ATF3 | B | POLR2A + EZH2 | 30.1 | |
| 22 | ATF4 | B | POLR2A | 600.3 | |
| 19 | ATF5 | B | POLR2A | 24.3 | |
| 1 | ATF6 | B | POLR2A | 20.7 | |
| 6 | ATF6B | B | POLR2A | 116.1 | |
| 12 | ATF7 | B | POLR2A | 32.8 | |
| 4 | ATOH1 | T | EZH2 | 17.1 | 8 |
| 10 | ATOH7 | T | POLR2A + EZH2 | 2.9 | |
| 21 | BACH1 | B | POLR2A | 17.9 | |
| 6 | BACH2 | B | POLR2A | 4.9 | |
| 9 | BARHL1 | T | EZH2 | 49.1 | 12 |
| 1 | BARHL2 | T | EZH2 | 84.0 | 12 |
| 9 | BARX1 | T | EZH2 | 47.0 | 8 |

| | | | | | |
|---|---|---|---|---|---|
| 11 | BARX2 | T | EZH2 | 143.4 | 6 |
| 14 | BATF | B | POLR2A | 10.3 | |
| 11 | BATF2 | B/T | POLR2A | 4.9 | |
| 1 | BATF3 | T | EZH2 | 4.4 | 2 |
| 3 | BBX | B | POLR2A | 22.7 | |
| 2 | BCL11A | T | POLR2A > EZH2 | 6.5 | |
| 14 | BCL11B | T | POLR2A + EZH2 | 8.9 | |
| 3 | BCL6 | B | POLR2A | 208.4 | |
| 17 | BCL6B | B | POLR2A > EZH2 | 15.0 | |
| 6 | BCLAF1 | B | POLR2A | 58.4 | |
| 20 | BHLH23 | T | EZH2 | 0.5 | 4 |
| 3 | BHLHE40 | B | POLR2A | 316.4 | |
| 7 | BHLHA15 | T | POLR2A | 192.4 | 5 |
| 8 | BHLHE22 | T | EZH2 | 14.8 | 8 |
| 20 | BMP2 | B | EZH2 + POLR2A | 13.6 | |
| 15 | BNC1 | T | EZH2 + POLR2A | 18.0 | |
| 9 | BNC2 | T | POLR2A + EZH2 | 22.5 | |
| 5 | BRD9 | B | POLR2A | 30.1 | |
| 11 | BSX | T | EZH2 | 0.8 | 8 |
| 14 | MIDEAS | B | POLR2A | 37.0 | |
| 1 | CAMTA1 | B | POLR2A | 38.0 | |
| 17 | CAMTA2 | B | POLR2A | 91.0 | |
| 1 | CASZ1 | B | POLR2A + EZH2 | 54.9 | |
| 6 | CDC5L | B | POLR2A | 30.0 | |
| 5 | CDX1 | T | EZH2 | 86.8 | 4 |
| 13 | CDX2 | T | EZH2 | 29.9 | 10 |
| 19 | CEBPA | B | POLR2A + EZH2 | 72.5 | |
| 20 | CEBPB | B | POLR2A | 263.0 | |
| 19 | CEBPG | B | POLR2A | 35.0 | |
| 2 | CEBPZ | B | POLR2A | 30.9 | |
| 19 | CIC | B | POLR2A | 73.6 | |
| 9 | CIZ1 | B | POLR2A | 67.9 | |
| 4 | CLOCK | B | POLR2A | 10.1 | |
| 16 | CRAMP1 | B | POLR2A | 29.5 | |
| 9 | CREB3 | B | POLR2A | 98.1 | |
| 11 | CREB3L1 | B | POLR2A | 31.4 | |
| 7 | CREB3L2 | B | POLR2A | 46.7 | |
| 19 | CREB3L3 | T | neither | 304.8 | |
| 1 | CREB3L4 | B | POLR2A | 26.5 | |
| 7 | CREB5 | B | neither | 7.5 | |
| 12 | CREBL2 | B | POLR2A | 40.8 | |
| 11 | CREBZF | B | POLR2A | 74.0 | |
| 10 | CREM | B | POLR2A | 23.7 | |
| 19 | CRX | T | neither | 0.1 | |
| 16 | CTCF | B | POLR2A | 40.3 | |

| | | | | | |
|---|---|---|---|---|---|
| 20 | CTCFL | T | POLR2A | 7.5 | |
| 7 | CUX1 | B | POLR2A | 27.3 | |
| 12 | CUX2 | T | POLR2A + EZH2 | 10.6 | |
| 18 | CXXC1 | B | POLR2A | 72.9 | |
| 13 | DACH1 | B | neither | 7.1 | |
| 23 | DACH2 | T | neither | 5.0 | |
| 19 | DBP | B | POLR2A | 33.4 | |
| 11 | DBX1 | T | EZH2 | 0.6 | 8 |
| 12 | DBX2 | T | EZH2 | 3.2 | 4 |
| 12 | DDIT3 | B | POLR2A | 80.0 | |
| 11 | DEAF1 | B | POLR2A | 87.0 | |
| 2 | DLX1 | T | EZH2 | 10.0 | 14 |
| 2 | DLX2 | T | EZH2 + POLR2A | 1.5 | |
| 17 | DLX3 | T | EZH2 | 65.4 | 9 |
| 17 | DLX4 | T | EZH2 | 3.0 | 10 |
| 7 | DLX5 | T | EZH2 | 34.7 | 8 |
| 7 | DLX6 | T | EZH2 | 9.7 | 11 |
| 1 | DMBX1 | T | neither | 2.0 | |
| 9 | DMRT1 | T | EZH2 | 45.4 | 12 |
| 9 | DMRT2 | T | EZH2 | 18.4 | 7 |
| 9 | DMRT3 | T | EZH2 | 14.0 | 16 |
| 9 | DMRTA1 | T | POLR2A | 3.3 | |
| 1 | DMRTA2 | T | EZH2 | 4.5 | 12 |
| 1 | DMRTB1 | T | neither | 122.4 | |
| 19 | DMRTC2 | T | neither | 66.3 | |
| 7 | DMTF1 | B | POLR2A | 74.1 | |
| 7 | DNAJC2 | B | POLR2A | 13.8 | |
| 19 | DPF1 | T | POLR2A > EZH2 | 30.6 | |
| 11 | DPF2 | B | POLR2A | 63.7 | |
| 14 | DPF3 | T | EZH2 | 16.2 | 5 |
| 19 | DPRX | T | neither | 1.0 | |
| 1 | DR1 | B | POLR2A | 18.0 | |
| 11 | DRAP1 | B | POLR2A | 233.4 | |
| 10 | DRGX | T | EZH2 | 1.8 | 12 |
| 19 | DUXA | T | neither | 1.0 | |
| 3 | DZIP1L | B | POLR2A | 8.4 | |
| 20 | E2F1 | B | POLR2A | 9.5 | |
| 1 | E2F2 | T | POLR2A | 12.8 | |
| 6 | E2F3 | B | POLR2A | 14.1 | |
| 16 | E2F4 | B | POLR2A | 97.9 | |
| 8 | E2F5 | B | POLR2A | 8.9 | |
| 2 | E2F6 | B | POLR2A | 12.6 | |
| 12 | E2F7 | T | POLR2A | 4.7 | |
| 11 | E2F8 | T | POLR2A + EZH2 | 3.2 | |
| 5 | EBF1 | B | POLRA2 | 36.2 | |

| | | | | | |
|---|---|---|---|---|---|
| 10 | EBF3 | T | EZH2 | 5.0 | 7 |
| 20 | EBF4 | B | EZH2 | 71.4 | 8 |
| 9 | EDF1 | B | POLR2A | 289.4 | |
| 5 | EGR1 | B | POLR2A | 402.0 | |
| 10 | EGR2 | B | POLR2A > EZH2 | 11.0 | |
| 2 | EGR4 | T | EZH2 | 14.8 | 9 |
| 11 | EHF | T | POLR2A | 147.3 | |
| 13 | ELF1 | B | POLR2A | 50.6 | |
| 4 | ELF2 | B | POLR2A | 24.9 | |
| 1 | ELF3 | T | POLR2A | 29.0 | |
| 23 | ELF4 | B | POLR2A | 25.2 | |
| 11 | ELF5 | T | POLR2A | 3.9 | |
| 23 | ELK1 | B | POLR2A | 46.2 | |
| 12 | ELK3 | B | POLR2A | 48.5 | |
| 1 | ELK4 | B | POLR2A | 15.7 | |
| 2 | EMX1 | T | EZH2 | 18.8 | 19 |
| 10 | EMX2 | T | EZH2 | 20.2 | 12 |
| 2 | EN1 | T | EZH2 | 9.4 | 12 |
| 7 | EN2 | T | EZH2 | 79.3 | 8 |
| 3 | EOMES | T | EZH2 | 26.9 | 12 |
| 2 | EPAS1 | B | POLR2A | 204.9 | |
| 19 | ERF | B | POLR2A | 72.8 | |
| 21 | ERG | B | EZH2 > POLR2A | 28.1 | |
| 6 | ESR1 | T | EZH2+POLR2A | 45.6 | |
| 14 | ESR2 | T | POLR2A + EZH2 | 1.1 | |
| 11 | ESRRA | B | POLR2A | 65.6 | |
| 14 | ESRRB | T | EZH2 | 11.8 | 8 |
| 1 | ESRRG | T | EZH2 | 3.7 | 8 |
| 23 | ESX1 | T | EZH2 | 19.8 | 4 |
| 11 | ETS1 | B | POLR2A | 48.2 | |
| 21 | ETS2 | B | POLR2A | 242.3 | |
| 7 | ETV1 | B | POLR2A | 174.6 | |
| 19 | ETV2 | T | POLR2A | 1.8 | |
| 1 | ETV3 | B | POLR2A | 13.3 | |
| 1 | ETV3L | T | neither | 0.8 | |
| 17 | ETV4 | B | POLR2A | 6.4 | |
| 3 | ETV5 | B | POLR2A | 17.1 | |
| 12 | ETV6 | B | POLR2A | 25.0 | |
| 6 | ETV7 | B | EZH2 + POLR2A | 5.0 | |
| 11 | PLEKHB1 | B | POLR2A > EZH2 | 207.9 | |
| 7 | EVX1 | T | EZH2 | 5.0 | 7 |
| 2 | EVX2 | T | EZH2 | 2.2 | 6 |
| 17 | EZH1 | B | POLR2A | 100.7 | |
| 7 | EZH2 | B | POLR2A | 14.8 | |
| 5 | FAM170A | T | neither | 36.8 | |

| | | | | | |
|---|---|---|---|---|---|
| 2 | FBXO41 | T | POLR2A | 60.1 | |
| 2 | FEV | T | EZH2 | 4.5 | 12 |
| 2 | FIGLA | T | EZH2 | 0.8 | 6 |
| 19 | FIZ1 | B | POLR2A | 9.4 | |
| 11 | FLI1 | B | EZH2 + POLR2A | 18.6 | |
| 14 | FOS | B | POLR2A | 429.6 | |
| 19 | FOSB | B | POLR2A | 20.5 | |
| 11 | FOSL1 | B | POLR2A + EZH2 | 12.2 | |
| 2 | FOSL2 | B | POLR2A | 144.2 | |
| 14 | FOXA1 | T | POLR2A > EZH2 | 21.2 | |
| 20 | FOXA2 | T | EZH2 > POLR2A | 14.2 | |
| 19 | FOXA3 | T | POLR2A | 21.1 | |
| 15 | FOXB1 | T | EZH2 | 3.4 | 8 |
| 9 | FOXB2 | T | EZH2 | 0.1 | 9 |
| 6 | FOXC1 | B | POLR2A + EZH2 | 63.9 | |
| 16 | FOXC2 | T | EZH2 > POLR2A | 36.0 | |
| 1 | FOXD2 | T | EZH2 | 16.5 | 5 |
| 1 | FOXD3 | T | EZH2 > POLR2A | 49.5 | |
| 9 | FOXD4 | T | EZH2 | 3.4 | 1 |
| 2 | FOXD4L1 | T | EZH2 | 1.5 | 3 |
| 9 | FOXD4L3 | T | neither | 0.2 | |
| 9 | FOXD4L5 | T | neither | 0.1 | |
| 9 | FOXD4L6 | T | EZH2 | 0.3 | 1 |
| 9 | FOXE1 | T | EZH2 | 164.5 | 11 |
| 1 | FOXE3 | T | EZH2 | 1.2 | 4 |
| 16 | FOXF1 | B | EZH2 | 49.3 | 8 |
| 6 | FOXF2 | T | EZH2 | 130.0 | 12 |
| 14 | FOXG1 | T | EZH2 | 25.2 | 4 |
| 8 | FOXH1 | T | EZH2 + POLR2A | 0.8 | |
| 5 | FOXI1 | T | neither | 32.2 | |
| 10 | FOXI2 | T | EZH2 | 2.0 | 1 |
| 17 | FOXJ1 | T | EZH2 > POLR2A | 15.7 | |
| 12 | FOXJ2 | B | POLR2A | 30.8 | |
| 1 | FOXJ3 | B | POLR2A | 31.9 | |
| 17 | FOXK2 | B | POLR2A | 19.6 | |
| 16 | FOXL1 | T | EZH2 | 17.9 | 4 |
| 3 | FOXL2 | T | EZH2 | 102.0 | 12 |
| 12 | FOXM1 | B | POLR2A | 15.8 | |
| 17 | FOXN1 | T | POLR2A | 64.3 | |
| 2 | FOXN2 | B | POLR2A | 11.6 | |
| 14 | FOXN3 | B | POLR2A | 16.5 | |
| 12 | FOXN4 | T | EZH2 | 23.6 | 8 |
| 13 | FOXO1 | B | POLR2A | 66.8 | |
| 6 | FOXO3 | B | POLR2A | 50.2 | |
| 23 | FOXO4 | B | POLR2A | 45.3 | |

| | | | | | |
|---|---|---|---|---|---|
| 1 | FOXO6 | B | POLR2A + EZH2 | 10.1 | |
| 3 | FOXP1 | B | POLR2A | 18.3 | |
| 7 | FOXP2 | T | EZH2 + POLR2A | 14.7 | |
| 23 | FOXP3 | B | POLR2A | 1.9 | |
| 6 | FOXP4 | B | POLR2A | 41.0 | |
| 6 | FOXQ1 | T | EZH2 + POLR2A | 36.0 | |
| 11 | FOXR1 | T | neither | 9.6 | |
| 23 | FOXR2 | T | neither | 2.6 | |
| 20 | FOXS1 | T | POLR2A | 63.8 | |
| 21 | GABPA | B | POLR2A | 17.8 | |
| 15 | GABPB1 | B | POLR2A | 11.1 | |
| 23 | GATA1 | T | POLR2A | 3.7 | |
| 3 | GATA2 | B | EZH2 > POLR2A | 55.9 | |
| 10 | GATA3 | T | EZH2 > POLR2A | 269.4 | |
| 8 | GATA4 | T | EZH2 | 144.1 | 15 |
| 20 | GATA5 | T | EZH2 | 13.9 | 7 |
| 18 | GATA6 | T | EZH2 | 54.7 | 11 |
| 7 | GBX1 | T | EZH2 | 15.4 | 13 |
| 2 | GBX2 | T | EZH2 | 2.4 | 4 |
| 6 | GCM2 | T | EZH2 | 1.4 | 6 |
| 1 | GFI1 | T | EZH2 | 9.7 | 4 |
| 9 | GFI1B | T | neither | 5.5 | |
| 12 | GLI1 | B | POLR2A | 5.9 | |
| 12 | GLI2 | B | POLR2A | 6.1 | |
| 12 | GLI3 | B | POLR2A | 15.2 | |
| 1 | GLIS1 | B | EZH2 | 5.7 | 9 |
| 2 | GLIS2 | B | EZH2 | 36.1 | 8 |
| 1 | GLIS3 | T | POLR2A + EZH2 | 4.8 | |
| 1 | GMEB1 | B | POLR2A | 7.0 | |
| 1 | GON4L | B | POLR2A | 15.0 | |
| 2 | GRHL1 | T | POLR2A | 160.8 | |
| 8 | GRHL2 | T | POLR2A | 41.4 | |
| 1 | GRHL3 | T | POLR2A + EZH2 | 141.8 | |
| 19 | GRLF1 | B | POLR2A | 26.8 | |
| 14 | GSC | T | EZH2 | 10.5 | 7 |
| 22 | GSC2 | T | EZH2 | 3.7 | 8 |
| 13 | GSX1 | T | EZH2 | 1.2 | 6 |
| 4 | GSX2 | T | EZH2 | 0.8 | 14 |
| 20 | GZF1 | B | POLR2A | 12.7 | |
| 5 | HAND1 | T | EZH2 | 122.8 | 16 |
| 4 | HAND2 | T | EZH2 | 86.4 | 10 |
| 7 | HBP1 | B | POLR2A | 41.2 | |
| 23 | HDX | B | POLR2A | 2.9 | |
| 4 | HELT | T | EZH2 | 1.9 | 4 |
| 4 | HERC5 | B | POLR2A+EZH2 | 26.8 | |

| | | | | | |
|---|---|---|---|---|---|
| 3 | HES1 | B | POLR2A | 250.4 | |
| 1 | HES2 | T | EZH2 + POLR2A | 28.9 | |
| 1 | HES3 | T | EZH2 | 0.4 | 6 |
| 1 | HES4 | B | EZH2 + POLR2A | 64.6 | |
| 1 | HES5 | T | EZH2 | 8.8 | 12 |
| 17 | HES7 | T | EZH2 | 7.0 | 6 |
| 3 | HESX1 | B | POLR2A | 2.1 | |
| 6 | HEY2 | B | EZH2 > POLR2A | 56.3 | |
| 1 | HEYL | B | EZH2 + POLR2A | 116.3 | |
| 10 | HHEX | B | EZH2 | 50.3 | 13 |
| 17 | HIC1 | B | EZH2 + POLR2A | 30.1 | |
| 22 | HIC2 | B | POLR2A | 5.7 | |
| 14 | HIF1A | B | POLR2A | 82.9 | |
| 19 | HIF3A | B | POLR2A | 64.9 | |
| 6 | HIVEP1 | B | POLR2A | 8.1 | |
| 6 | HIVEP2 | B | POLR2A | 11.1 | |
| 1 | HIVEP3 | B | POLR2A | 6.4 | |
| 19 | ZNF875 | B | POLR2A | 25.8 | |
| 17 | HLF | B | POLR2A + EZH2 | 49.2 | |
| 1 | HLX | B | EZH2 | 32.3 | 7 |
| 8 | HMBOX1 | B | POLR2A | 8.2 | |
| 4 | HMX1 | T | EZH2 | 2.3 | 4 |
| 10 | HMX2 | T | EZH2 | 1.6 | 11 |
| 10 | HMX3 | T | EZH2 | 1.4 | 7 |
| 12 | HNF1A | T | POLR2A | 8.7 | |
| 17 | HNF1B | T | EZH2 + POLR2A | 90.4 | |
| 8 | HNF4G | T | neither | 24.4 | |
| 14 | HOMEZ | B | POLR2A | 9.7 | |
| 4 | HOPX | T | EZH2 + POLR2A | 746.5 | |
| 7 | HOXA1 | T | EZH2 > POLR2A | 3.6 | |
| 7 | HOXA10 | T | EZH2 > POLR2A | 70.5 | |
| 7 | HOXA11 | T | EZH2=POLR2A | 200.7 | 8 |
| 7 | HOXA13 | T | EZH2 + POLR2A | 71.9 | |
| 7 | HOXA2 | T | EZH2 > POLR2A | 8.1 | |
| 7 | HOXA3 | T | EZH2 | 14.4 | 10 |
| 7 | HOXA4 | T | EZH2 | 44.6 | 3 |
| 7 | HOXA5 | T | EZH2 + POLR2A | 117.9 | |
| 7 | HOXA6 | T | EZH2 + POLR2A | 14.8 | |
| 7 | HOXA7 | T | EZH2 | 3.1 | 9 |
| 7 | HOXA9 | T | EZH2 | 33.6 | 11 |
| 17 | HOXB1 | T | EZH2 | 0.8 | |
| 17 | HOXB13 | T | EZH2 | 118.8 | 16 |
| 17 | HOXB2 | B | EZH2 + POLR2A | 47.2 | |
| 17 | HOXB3 | T | EZH2 > POLR2A | 33.2 | |
| 17 | HOXB4 | B/T | EZH2 | 22.9 | 3 |

| | | | | | |
|---|---|---|---|---|---|
| 17 | HOXB5 | T | EZH2 > POLR2A | 34.5 | |
| 17 | HOXB6 | T | EZH2 | 40.1 | 11 |
| 17 | HOXB7 | T | EZH2 > POLR2A | 36.9 | |
| 17 | HOXB8 | T | EZH2 + POLR2A | 37.3 | |
| 17 | HOXB9 | T | EZH2 + POLR2A | 35.4 | |
| 12 | HOXC10 | T | EZH2 | 61.6 | 7 |
| 12 | HOXC11 | T | EZH2 | 6.7 | 8 |
| 12 | HOXC12 | T | EZH2 | 5.4 | 13 |
| 12 | HOXC13 | T | EZH2 | 12.9 | 8 |
| 12 | HOXC4 | T | EZH2 | 42.8 | 19 |
| 12 | HOXC5 | T | EZH2 | 17.8 | 19 |
| 12 | HOXC6 | T | EZH2 | 37.9 | 19 |
| 12 | HOXC8 | T | EZH2 | 22.2 | 15 |
| 12 | HOXC9 | T | EZH2 | 22.3 | 22 |
| 2 | HOXD1 | T | EZH2 > POLR2A | 9.1 | |
| 2 | HOXD10 | T | EZH2 | 103.7 | 6 |
| 2 | HOXD11 | T | EZH2 | 15.5 | 6 |
| 2 | HOXD12 | T | EZH2 | 0.6 | 8 |
| 2 | HOXD13 | T | EZH2 | 34.0 | 11 |
| 2 | HOXD3 | T | EZH2 | 29.2 | 14 |
| 2 | HOXD4 | T | EZH2 | 36.1 | 8 |
| 2 | HOXD8 | T | EZH2 | 50.5 | 12 |
| 2 | HOXD9 | T | EZH2 > POLR2A | 173.5 | |
| 8 | HSF1 | B | POLR2A | 108.6 | |
| 6 | HSF2 | B | POLR2A | 33.0 | |
| 16 | HSF4 | B | EZH2 + POLR2A | 108.1 | |
| 17 | HSF5 | T | neither | 64.5 | |
| 20 | ID1 | B | POLR2A | 204.6 | |
| 2 | ID2 | B | POLR2A | 126.5 | |
| 1 | ID3 | B | POLR2A | 357.9 | |
| 1 | IFI16 | B | POLR2A | 112.3 | |
| 7 | IKZF1 | T | POLR2A > EZH2 | 34.7 | |
| 2 | IKZF2 | B | POLR2A | 9.0 | |
| 17 | IKZF3 | T | POLR2A + EZH2 | 36.8 | |
| 12 | IKZF4 | B | POLR2A | 10.8 | |
| 10 | IKZF5 | B | POLR2A | 25.5 | |
| 20 | INSM1 | T | EZH2 | 53.3 | 7 |
| 14 | INSM2 | T | EZH2 | 2.4 | 11 |
| 5 | IRF1 | B | POLR2A | 42.2 | |
| 4 | IRF2 | B | POLR2A | 28.9* | |
| 19 | IRF3 | B | POLR2A | 86.9 | |
| 6 | IRF4 | T | POLR2A | 33.4 | |
| 7 | IRF5 | B | POLR2A | 11.8 | |
| 1 | IRF6 | T | POLR2A | 188.7 | |
| 11 | IRF7 | B | POLR2A | 44.3 | |

| | | | | | |
|---|---|---|---|---|---|
| 16 | IRF8 | B | POLR2A | 21.0 | |
| 14 | IRF9 | B | POLR2A | 96.0 | |
| 5 | IRX1 | T | EZH2 | 36.0 | 8 |
| 5 | IRX2 | T | POLR2A + EZH2 | 40.2 | |
| 16 | IRX3 | T | EZH2 | 31.9 | 4 |
| 5 | IRX4 | T | EZH2 + POLR2A | 31.6 | |
| 16 | IRX5 | T | EZH2 > POLR2A | 29.1 | |
| 16 | IRX6 | T | EZH2 > POLR2A | 13.4 | |
| 5 | ISL1 | T | EZH2 | 28.0 | 6 |
| 15 | ISL2 | T | EZH2 + POLR2A | 5.9 | |
| 22 | ISX | T | POLR2A | 24.8 | |
| 7 | JAZF1 | B | POLR2A | 50.2 | |
| 14 | JDP2 | B | POLR2A | 27.6 | |
| 1 | JUN | B | POLR2A | 333.6 | |
| 19 | JUNB | B | POLR2A | 788.0 | |
| 19 | JUND | B | POLR2A | 305.5 | |
| 7 | KIAA1549 | B | POLR2A > EZH2 | 3.2 | |
| 19 | KLF1 | T | POLR2A | 1.4 | |
| 8 | KLF10 | B | POLR2A | 57.4 | |
| 2 | KLF11 | B | POLR2A | 27.1 | |
| 13 | KLF12 | B | POLR2A | 10.0 | |
| 15 | KLF13 | B | POLR2A | 44.9 | |
| 7 | KLF14 | T | EZH2 | 2.4 | 14 |
| 3 | KLF15 | B | EZH2 + POLR2A | 54.8 | |
| 19 | KLF16 | B | POLR2A | 27.3 | |
| 1 | KLF17 | T | neither | 24.5 | |
| 19 | KLF2 | B | POLR2A + EZH2 | 106.7 | |
| 4 | KLF3 | B | POLR2A | 39.5 | |
| 13 | KLF5 | B | POLR2A | 77.5 | |
| 10 | KLF6 | B | POLR2A | 95.8 | |
| 2 | KLF7 | B | POLR2A | 18.1 | |
| 23 | KLF8 | B | POLR2A | 20.3 | |
| 9 | KLF9 | B | POLR2A | 122.3 | |
| 10 | LBX1 | T | EZH2 | 9.7 | 15 |
| 2 | LBX2 | T | EZH2 + POLR2A | 1.3 | |
| 10 | LCOR | B | neither | 6.1 | |
| 4 | LCORL | B | POLR2A | 2.7 | |
| 4 | LEF1 | T | EZH2 | 12.7 | 9 |
| 19 | LEUTX | T | neither | 0.1 | |
| 17 | LHX1 | T | EZH2 | 55.1 | 19 |
| 9 | LHX2 | T | EZH2 | 50.5 | 16 |
| 9 | LHX3 | T | EZH2 | 89.7 | 4 |
| 1 | LHX4 | B | EZH2 | 3.1 | 11 |
| 12 | LHX5 | T | EZH2 | 4.1 | 8 |
| 9 | LHX6 | T | EZH2 | 7.4 | 7 |

| | | | | | |
|---|---|---|---|---|---|
| 1 | LHX8 | T | EZH2 | 4.0 | 12 |
| 1 | LHX9 | T | EZH2 | 4.1 | 18 |
| 1 | LMX1A | T | EZH2 | 2.4 | 8 |
| 9 | LMX1B | T | EZH2 | 5.1 | 12 |
| 1 | MAEL | T | neither | 174.2 | |
| 16 | MAF | B | POLR2A + EZH2 | 76.4 | |
| 8 | MAFA | T | EZH2 | 16.5 | 12 |
| 20 | MAFB | T | EZH2 | 83.0 | 7 |
| 22 | MAFF | B | POLR2A | 71.3 | |
| 17 | MAFG | B | POLR2A | 30.4 | |
| 7 | MAFK | B | POLR2A | 89.6 | |
| 5 | MATR3 | B | POLR2A | 14.9 | |
| 14 | MAX | B | POLR2A | 54.2 | |
| 16 | MAZ | B | POLR2A | 35.6 | |
| 15 | MEF2A | B | POLR2A | 46.4 | |
| 19 | MEF2B | B | POLR2A | 18.6 | |
| 5 | MEF2C | B | POLR2A | 23.1 | |
| 1 | MEF2D | B | POLR2A | 75.2 | |
| 2 | MEIS1 | B | EZH2 + POLR2A | 43.6 | |
| 15 | MEIS2 | B | POLR2A > EZH2 | 33.7 | |
| 19 | MEIS3 | T | EZH2 + POLR2A | 40.1 | |
| 7 | MEOX2 | T | EZH2 | 54.2 | 4 |
| 15 | MESP2 | B | EZH2 + POLR2A | 1.0 | |
| 15 | MGA | B | POLR2A | 10.5 | |
| 1 | MIER1 | B | POLR2A | 20.9 | |
| 19 | MIER2 | B | POLR2A | 13.6 | |
| 5 | MIER3 | B | POLR2A | 15.0 | |
| 1 | MIXL1 | T | EZH2 | 2.3 | 2 |
| 11 | MIZF | B | POLR2A | 11.7 | |
| 15 | MKRN3 | T | POLR2A | 3.4 | |
| 10 | MKX | T | EZH2 | 16.2 | 10 |
| 19 | MLLT1 | B | POLR2A | 83.6 | |
| 9 | MLLT3 | B | POLR2A + EZH2 | 8.2 | |
| 17 | MLX | B | POLR2A | 58.5 | |
| 12 | MLXIP | B | POLR2A | 49.2 | |
| 7 | MLXIPL | B | EZH2 + POLR2A | 44.8 | |
| 17 | MNT | B | POLR2A | 21.1 | |
| 7 | MNX1 | T | EZH2 | 7.7 | 12 |
| 8 | MSC | B | EZH2 + POLR2A | 13.7 | |
| 4 | MSX1 | B | EZH2 + POLR2A | 14.2 | |
| 5 | MSX2 | T | EZH2 | 11.3 | 4 |
| 1 | MTF1 | B | POLR2A | 8.5 | |
| 2 | MXD1 | B | POLR2A | 19.2 | |
| 5 | MXD3 | B | POLR2A | 12.3 | |
| 4 | MXD4 | B | POLR2A | 67.0 | |

| | | | | | |
|---|---|---|---|---|---|
| 10 | MXI1 | B | POLR2A | 59.6 | |
| 6 | MYB | T | POLR2A + EZH2 | 12.6 | |
| 8 | MYBL1 | B | POLR2A | 7.6 | |
| 20 | MYBL2 | T | POLR2A | 37.9 | |
| 8 | MYC | B | POLR2A | 189.2 | |
| 1 | MYCL1 | T | POLR2A > EZH2 | 38.4 | |
| 2 | MYCN | T | EZH2 > POLR2A | 7.5 | |
| 12 | MYF5 | T | neither | 2.1 | |
| 12 | MYF6 | T | POLR2A | 255.0 | |
| 3 | MYNN | B | POLR2A | 10.1 | |
| 11 | MYOD1 | T | EZH2 | 21.7 | 6 |
| 1 | MYOG | T | neither | 58.0 | |
| 20 | MYT1 | T | EZH2 | 64.7 | 4 |
| 2 | MYT1L | T | neither | 37.0 | |
| 19 | MZF1 | B | POLR2A | 42.6 | |
| 12 | NANOG | T | neither | 0.2 | |
| 2 | NCOA1 | B | POLR2A | 26.8 | |
| 20 | NCOA3 | B | POLR2A | 23.2 | |
| 2 | NEUROD1 | T | EZH2 | 306.4 | 7 |
| 12 | NEUROD4 | T | POLR2A | 15.6 | |
| 7 | NEUROD6 | T | EZH2 | 38.3 | 5 |
| 5 | NEUROG1 | T | EZH2 | 0.2 | 8 |
| 4 | NEUROG2 | T | EZH2 | 3.8 | 5 |
| 10 | NEUROG3 | T | EZH2 | 1.6 | 6 |
| 16 | NFAT5 | B | POLR2A | 15.2 | |
| 18 | NFATC1 | B | POLR2A + EZH2 | 10.8 | |
| 16 | NFATC3 | B | POLR2A | 13.8 | |
| 14 | NFATC4 | B | POLR2A + EZH2 | 49.4 | |
| 12 | NFE2 | T | POLR2A | 47.4 | |
| 17 | NFE2L1 | B | POLR2A | 224.8 | |
| 2 | NFE2L2 | B | POLR2A | 97.7 | |
| 7 | NFE2L3 | B | POLR2A | 11.2 | |
| 1 | NFIA | B | POLR2A | 18.5 | |
| 9 | NFIB | B | POLR2A + EZH2 | 26.4 | |
| 19 | NFIC | B | POLR2A | 70.1 | |
| 9 | NFIL3 | B | POLR2A | 125.1 | |
| 19 | NFIX | B | EZH2 > POLR2A | 82.1 | |
| 4 | NFKB1 | B | POLR2A | 24.8 | |
| 10 | NFKB2 | B | POLR2A | 44.1 | |
| 11 | NFRKB | B | POLR2A | 25.5 | |
| 9 | NFX1 | B | POLR2A | 24.3 | |
| 4 | NFXL1 | B | POLR2A | 8.5 | |
| 6 | NFYA | B | POLR2A | 23.5 | |
| 12 | NFYB | B | POLR2A | 50.8 | |
| 1 | NFYC | B | POLR2A | 37.5 | |

| | | | | | |
|---|---|---|---|---|---|
| 1 | NHLH1 | T | POLR2A | 2.3 | |
| 1 | NHLH2 | T | EZH2 | 7.3 | 7 |
| 23 | NKRF | B | POLR2A | 9.8 | |
| 14 | NKX2-1 | T | EZH2 | 352.6 | 8 |
| 20 | NKX2-2 | T | EZH2 | 43.8 | 12 |
| 10 | NKX2-3 | T | EZH2 | 71.4 | 10 |
| 20 | NKX2-4 | T | EZH2 | 3.2 | 8 |
| 5 | NKX2-5 | T | EZH2 | 113.7 | 11 |
| 8 | NKX2-6 | T | EZH2 | 1.4 | 4 |
| 14 | NKX2-8 | T | EZH2 | 17.6 | 4 |
| 14 | NKX3-1 | T | EZH2 > POLR2A | 250.5 | |
| 4 | NKX3-2 | T | EZH2 | 33.0 | 15 |
| 4 | NKX6-1 | T | EZH2 | 125.7 | 10 |
| 10 | NKX6-2 | T | EZH2 | 147.1 | 3 |
| 8 | NKX6-3 | T | EZH2 | 12.7 | 2 |
| 7 | NOBOX | T | neither | 0.9 | |
| 2 | NOTO | T | EZH2 | 0.2 | 6 |
| 19 | NPAS1 | T | EZH2 | 10.8 | 12 |
| 2 | NPAS2 | B | POLR2A | 26.6 | |
| 11 | NPAS4 | B | EZH2 + POLR2A | 9.1 | |
| 23 | NR0B1 | T | EZH2 | 42.5 | 3 |
| 17 | NR1D1 | B | POLR2A | 115.4 | |
| 3 | NR1D2 | B | POLR2A | 33.7 | |
| 19 | NR1H2 | B | POLR2A | 110.4 | |
| 11 | NR1H3 | B | POLR2A | 27.5 | |
| 12 | NR1H4 | T | POLR2A | 39.1 | |
| 3 | NR1I2 | T | POLR2A | 42.6 | |
| 1 | NR1I3 | T | neither | 98.7 | |
| 12 | NR2C1 | B | POLR2A | 16.5 | |
| 3 | NR2C2 | B | POLR2A | 21.3 | |
| 6 | NR2E1 | T | EZH2 | 20.0 | 13 |
| 15 | NR2E3 | T | neither | 0.6 | |
| 5 | NR2F1 | B | EZH2 | 64.7 | 12 |
| 15 | NR2F2 | B | EZH2 | 66.3 | 11 |
| 19 | NR2F6 | B | POLR2A | 39.0 | |
| 5 | NR3C1 | B | POLR2A | 24.0 | |
| 4 | NR3C2 | B | POLR2A + EZH2 | 9.6 | |
| 12 | NR4A1 | B | POLR2A | 309.2 | |
| 2 | NR4A2 | B | POLR2A > EZH2 | 26.0 | |
| 9 | NR4A3 | B/T | POLR2A + EZH2 | 10.0 | |
| 9 | NR5A1 | T | EZH2 | 221.6 | 8 |
| 1 | NR5A2 | T | EZH2 + POLR2A | 7.8 | |
| 9 | NR6A1 | B | POLR2A | 3.9 | |
| 7 | NRF1 | B | POLR2A | 15.6 | |
| 14 | NRL | B | POLR2A + EZH2 | 2.1 | |

| | | | | | |
|---|---|---|---|---|---|
| 4 | NSD2 | B | POLR2A | 11.0 | |
| 21 | OLIG2 | T | EZH2 | 34.9 | 12 |
| 6 | OLIG3 | T | EZH2 | 0.1 | 12 |
| 15 | ONECUT1 | T | EZH2 | 5.5 | 8 |
| 18 | ONECUT2 | T | EZH2 | 4.5 | 7 |
| 19 | ONECUT3 | T | EZH2 | 2.6 | 4 |
| 2 | OSR1 | T | EZH2 | 154.1 | 11 |
| 8 | OSR2 | T | EZH2 | 246.1 | 3 |
| 5 | OTP | T | EZH2 | 9.4 | 15 |
| 2 | OTX1 | T | EZH2 + POLR2A | 28.1 | |
| 14 | OTX2 | T | EZH2 | 24.4 | 10 |
| 11 | OVOL1 | T | EZH2 | 43.5 | 10 |
| 20 | OVOL2 | T | EZH2 | 15.6 | 12 |
| 22 | PATZ1 | B | POLR2A | 24.4 | |
| 12 | PAWR | B | POLR2A | 26.9 | |
| 20 | PAX1 | T | EZH2 | 0.6 | 8 |
| 10 | PAX2 | T | EZH2 | 52.8 | 19 |
| 2 | PAX3 | T | EZH2 | 4.6 | 10 |
| 7 | PAX4 | T | EZH2 | 1.3 | 3 |
| 9 | PAX5 | T | EZH2 | 17.6 | 4 |
| 11 | PAX6 | T | EZH2 | 36.9 | 15 |
| 1 | PAX7 | T | EZH2 | 1.7 | 16 |
| 2 | PAX8 | T | POLR2A > EZH2 | 1193.0 | |
| 14 | PAX9 | T | EZH2 | 81.1 | 4 |
| 1 | PBX1 | B | POLR2A | 45.9 | |
| 6 | PBX2 | B | POLR2A | 87.8 | |
| 9 | PBX3 | B | POLR2A | 37.9 | |
| 19 | PBX4 | B | POLR2A | 7.6 | |
| 10 | PCGF6 | B | POLR2A | 13.3 | |
| 13 | PDX1 | T | EZH2 | 9.2 | 15 |
| 11 | PGR | T | EZH2 | 111.0 | 12 |
| 12 | PHB2 | B | POLR2A | 241.9 | |
| 20 | PHF20 | B | POLR2A | 21.0 | |
| 11 | PHOX2A | T | EZH2 > POLR2A | 2.7 | |
| 4 | PHOX2B | T | EZH2 | 4.1 | 4 |
| 5 | PITX1 | T | EZH2 | 816.4 | 14 |
| 4 | PITX2 | T | EZH2 | 24.5 | 12 |
| 10 | PITX3 | T | EZH2 | 9.3 | 7 |
| 21 | PKNOX1 | B | POLR2A | 17.4 | |
| 11 | PKNOX2 | B | POLR2A + EZH2 | 11.0 | |
| 8 | PLAG1 | B/T | EZH2 | 4.2 | 5 |
| 6 | PLAGL1 | B | POLR2A + EZH2 | 36.5 | |
| 20 | PLAGL2 | B | POLR2A | 12.7 | |
| 2 | PLEK | B | POLR2A | 17.6 | |
| 3 | POU1F1 | T | neither | 111.3 | |

| | | | | | |
|---|---|---|---|---|---|
| 1 | POU2F1 | B | POLR2A | 8.2 | |
| 19 | POU2F2 | B | POLR2A | 6.4 | |
| 11 | POU2F3 | T | POLR2A > EZH2 | 102.4 | |
| 1 | POU3F1 | T | EZH2 | 34.0 | 12 |
| 6 | POU3F2 | T | EZH2 | 10.7 | 11 |
| 2 | POU3F3 | T | EZH2 | 77.4 | 10 |
| 23 | POU3F4 | T | EZH2 | 28.9 | 2 |
| 13 | POU4F1 | T | EZH2 | 2.1 | 10 |
| 4 | POU4F2 | T | EZH2 | 2.8 | 4 |
| 5 | POU4F3 | T | EZH2 | 0.1 | 5 |
| 6 | POU5F1 | B | POLR2A | 6.3 | |
| 12 | POU6F1 | B | POLR2A | 24.0 | |
| 7 | POU6F2 | T | EZH2 | 1.9 | 4 |
| 6 | PPARD | B | POLR2A | 43.5 | |
| 3 | PPARG | B | POLR2A | 11.0 | |
| 6 | PRDM1 | B | POLR2A | 19.6 | |
| 11 | PRDM10 | B | POLR2A | 7.0 | |
| 9 | PRDM12 | T | EZH2 | 0.8 | 15 |
| 6 | PRDM13 | T | EZH2 | 0.4 | 10 |
| 8 | PRDM14 | T | EZH2 | 0.9 | 12 |
| 21 | PRDM15 | B | POLR2A | 5.9 | |
| 1 | PRDM16 | T | EZH2 | 20.5 | 6 |
| 1 | PRDM2 | B | POLR2A | 26.5 | |
| 12 | PRDM4 | B | POLR2A | 23.2 | |
| 4 | PRDM5 | B | POLR2A | 5.3 | |
| 5 | PRDM6 | T | EZH2 | 35.5 | 16 |
| 16 | PRDM7 | T | POLR2A | 4.0 | |
| 4 | PRDM8 | T/B | EZH2 | 25.7 | 12 |
| 5 | PRDM9 | T | POLR2A- | 6.9 | |
| 2 | PREB | B | POLR2A | 92.7 | |
| 5 | PROP1 | T | EZH2 | 2.0 | 3 |
| 1 | PROX1 | T | EZH2 + POLR2A | 30.5 | |
| 14 | PROX2 | B | neither | 24.7 | |
| 1 | PRRX1 | T | EZH2 | 88.5 | 12 |
| 9 | PRRX2 | T | EZH2 | 46.7 | 4 |
| 10 | PTF1A | T | EZH2 | 30.3 | 4 |
| 17 | RARA | B | POLR2A | 42.6 | |
| 3 | RARB | B | POLR2A | 12.7 | |
| 12 | RARG | B | POLR2A | 36.4 | |
| 18 | RAX | T | EZH2 | 0.3 | 15 |
| 19 | RAX2 | T | neither | 1.9 | |
| 7 | RBAK | B | POLR2A | 8.8 | |
| 4 | RBPJ | B | POLR2A | 35.0 | |
| 20 | RBPJL | T | EZH2 | 666.8 | 4 |
| 9 | RC3H2 | B | POLR2A | 14.3 | |

| | | | | | |
|---|---|---|---|---|---|
| 14 | RCOR1 | B | POLR2A | 28.2 | |
| 11 | RCOR2 | T | POLR2A + EZH2 | 13.2 | |
| 1 | RCOR3 | B | POLR2A | 29.8 | |
| 2 | REL | B | POLR2A | 8.3 | |
| 11 | RELA | B | POLR2A | 88.4 | |
| 19 | RELB | B | POLR2A | 15.0 | |
| 1 | RERE | B | POLR2A | 79.1 | |
| 4 | REST | B | POLR2A | 13.1 | |
| 19 | RFX1 | B | POLR2A | 22.6 | |
| 19 | RFX2 | B | POLR2A | 19.1 | |
| 9 | RFX3 | B | POLR2A | 8.3 | |
| 12 | RFX4 | T | EZH2 | 92.0 | 12 |
| 1 | RFX5 | B | POLR2A | 30.9 | |
| 6 | RFX6 | T | EZH2 | 1.8 | 8 |
| 19 | RFXANK | B | POLR2A | 48.2 | |
| 23 | RHOXF1 | B | neither | 5.3 | |
| 23 | RHOXF2 | T | neither | 5.5 | |
| 1 | RLF | B | POLR2A | 20.8 | |
| 15 | RORA | B | EZH2 + POLR2A | 30.6 | |
| 9 | RORB | T | EZH2 | 17.0 | 12 |
| 1 | RORC | T | POLR2A | 47.7 | |
| 6 | RREB1 | B | POLR2A | 27.2 | |
| 21 | RUNX1 | B | POLR2A | 13.1 | |
| 6 | RUNX2 | B | POLR2A | 9.9 | |
| 1 | RUNX3 | B | POLR2A | 14.2 | |
| 6 | RXRB | B | POLR2A | 74.6 | |
| 1 | RXRG | T | EZH2 | 43.1 | 8 |
| 16 | SALL1 | T | EZH2 | 50.8 | 4 |
| 14 | SALL2 | B | POLR2A | 88.2 | |
| 18 | SALL3 | T | EZH2 | 8.8 | 3 |
| 20 | SALL4 | T | POLR2A | 11.9 | |
| 3 | SATB1 | B | EZH2 + POLR2A | 24.2 | |
| 2 | SATB2 | T | EZH2 + POLR2A | 4.4 | 10 |
| 8 | SCRT1 | T | EZH2 | 140.7 | 4 |
| 20 | SCRT2 | T | EZH2 | 6.9 | 6 |
| 17 | SEBOX | T | POLR2A | 0.3 | |
| 3 | SHOX2 | T | EZH2 | 26.1 | 16 |
| 6 | SIM1 | T | EZH2 | 5.8 | 14 |
| 21 | SIM2 | T | EZH2 | 40.7 | 10 |
| 14 | SIX1 | T | EZH2 | 30.5 | 10 |
| 2 | SIX2 | T | EZH2 + POLR2A | 22.7 | |
| 2 | SIX3 | T | EZH2 | 52.0 | 15 |
| 14 | SIX4 | B | POLR2A | 5.4 | |
| 19 | SIX5 | B | POLR2A | 41.3 | |
| 14 | SIX6 | T | EZH2 | 55.2 | 8 |

| | | | | | |
|---|---|---|---|---|---|
| 1 | SKI | B | POLR2A | 104.4 | |
| 3 | SKIL | B | POLR2A | 20.0 | |
| 15 | SKOR1 | T | EZH2 | 13.6 | 14 |
| 4 | SMAD1 | B | POLR2A | 12.8 | |
| 18 | SMAD2 | B | POLR2A | 6.8 | |
| 15 | SMAD3 | B | POLR2A | 55.2 | |
| 18 | SMAD4 | B | POLR2A | 32.3 | |
| 5 | SMAD5 | B | POLR2A | 26.8 | |
| 13 | SMAD9 | B | EZH2 + POLR2A | 16.9 | |
| 20 | SNAI1 | B | POLR2A | 16.3 | |
| 8 | SNAI2 | B | POLR2A>EZH2 | 98.7 | |
| 16 | SNAI3 | B | POLR2A | 4.8 | |
| 9 | SNAPC4 | B | POLR2A | 24.0 | |
| 9 | SOHLH1 | T | EZH2 | 40.2 | 1 |
| 13 | SOX1 | T | EZH2 | 5.2 | 1 |
| 22 | SOX10 | T | POLR2A + EZH2 | 0.9 | |
| 2 | SOX11 | T | EZH2 | 2.3 | 7 |
| 20 | SOX12 | B | POLR2A | 20.6 | |
| 1 | SOX13 | B | POLR2A | 54.1 | |
| 3 | SOX14 | T | EZH2 | 0.6 | 12 |
| 17 | SOX15 | B | POLR2A | 51.6 | |
| 8 | SOX17 | T | EZH2 > POLR2A | 47.3 | |
| 20 | SOX18 | B | EZH2 + POLR2A | 51.7 | |
| 3 | SOX2 | T | EZH2 + POLR2A | 68.0 | |
| 13 | SOX21 | T | EZH2 | 27.1 | 8 |
| 23 | SOX3 | T | EZH2 + POLR2A | 5.6 | |
| 5 | SOX30 | T | POLR2A- | 137.0 | |
| 6 | SOX4 | B | POLR2A > EZH2 | 26.8 | |
| 12 | SOX5 | B | POLR2A | 6.4 | |
| 11 | SOX6 | B | POLR2A | 8.0 | |
| 8 | SOX7 | B | EZH2 + POLR2A | 34.2 | |
| 16 | SOX8 | T | EZH2 | 124.9 | 3 |
| 17 | SOX9 | B | POLR2A > EZH2 | 33.2 | |
| 12 | SP1 | B | POLR2A | 45.3 | |
| 2 | SP100 | B | POLR2A | 48.5 | |
| 2 | SP110 | B | POLR2A | 19.7 | |
| 17 | SP2 | B | POLR2A | 22.4 | |
| 2 | SP3 | B | POLR2A | 36.8 | |
| 7 | SP4 | B | POLR2A | 7.7 | |
| 2 | SP5 | T | POLR2A + EZH2 | 27.7 | |
| 17 | SP6 | T | EZH2 | 22.5 | 8 |
| 12 | SP7 | T | EZH2 | 1.6 | 4 |
| 7 | SP8 | T | EZH2 | 4.9 | 6 |
| 6 | SPDEF | T | POLR2A | 211.8 | |
| 11 | SPI1 | B | POLR2A | 27.0 | |

| | | | | | |
|---|---|---|---|---|---|
| 19 | SPIB | T | POLR2A + EZH2 | 21.9 | |
| 12 | SPIC | T | neither | 23.3 | |
| 17 | SREBF1 | B | POLR2A | 115.4 | |
| 22 | SREBF2 | B | POLR2A | 90.1 | |
| 6 | SRF | B | POLR2A | 99.7 | |
| 24 | SRY | T | neither | 4.5 | |
| 8 | ST18 | T | neither | 33.2 | |
| 2 | STAT1 | B | POLR2A | 45.1 | |
| 12 | STAT2 | B | POLR2A | 109.9 | |
| 17 | STAT3 | B | POLR2A | 101.4 | |
| 2 | STAT4 | B | POLR2A | 24.0 | |
| 17 | STAT5A | B | POLR2A | 47.3 | |
| 17 | STAT5B | B | POLR2A | 72.2 | |
| 12 | STAT6 | B | POLR2A | 161.8 | |
| 17 | SUZ12 | B | POLR2A | 20.4 | |
| 1 | TAL1 | T/B | EZH2 > POLR2A | 11.7 | |
| 9 | TAL2 | B | neither | 0.7 | |
| 7 | TAX1BP1 | B | POLR2A | 34.3 | |
| 2 | TBR1 | T | EZH2 | 42.0 | 14 |
| 22 | TBX1 | T | EZH2 | 34.8 | 12 |
| 11 | TBX10 | T | POLR2A | 5.2 | |
| 1 | TBX15 | T | EZH2 | 77.9 | 9 |
| 6 | TBX18 | T | EZH2 | 46.7 | 7 |
| 1 | TBX19 | B | POLR2A | 17.7 | |
| 17 | TBX2 | B | EZH2 > POLR2A | 130.4 | |
| 7 | TBX20 | T | EZH2 | 20.7 | 8 |
| 17 | TBX21 | T | EZH2 + POLR2A | 18.5 | |
| 23 | TBX22 | T | EZH2 | 38.6 | 3 |
| 12 | TBX3 | B | EZH2 > POLR2A | 62.3 | |
| 17 | TBX4 | T | EZH2 | 54.1 | 15 |
| 12 | TBX5 | T | EZH2 | 71.0 | 8 |
| 16 | TBX6 | B | POLR2A | 3.8 | |
| 6 | TBXT | T | EZH2 | 1.4 | 10 |
| 15 | TCF12 | B | POLR2A | 26.0 | |
| 20 | TCF15 | T | EZH2 > POLR2A | 11.7 | |
| 6 | TCF19 | B | POLR2A | 15.8 | |
| 22 | TCF20 | B | neither | 18.6 | |
| 6 | TCF21 | T | EZH2 | 73.6 | 10 |
| 2 | TCF23 | T | POLR2A | 103.4 | |
| 8 | TCF24 | T | EZH2 | 2.8 | 15 |
| 16 | TCF25 | B | POLR2A | 75.0 | |
| 19 | TCF3 | B | POLR2A | 36.3 | |
| 18 | TCF4 | B | POLR2A | 24.0 | |
| 5 | TCF7 | B | POLR2A | 12.1 | |
| 2 | TCF7L1 | B | POLR2A + EZH2 | 38.9 | |

| | | | | | |
|---|---|---|---|---|---|
| 10 | TCF7L2 | B | POLR2A | 26.1 | |
| 20 | TCFL5 | B | POLR2A | 22.7 | |
| 19 | TEAD2 | B | POLR2A | 32.7 | |
| 6 | TEAD3 | B | POLR2A | 108.9 | |
| 12 | TEAD4 | B | POLR2A | 12.6 | |
| 22 | TEF | B | POLR2A | 38.3 | |
| 16 | TERB1 | T | POLR2A | 17.4 | |
| 10 | TFAM | B | POLR2A | 12.0 | |
| 6 | TFAP2A | T | EZH2 + POLR2A | 63.2 | |
| 6 | TFAP2B | T | EZH2 | 17.4 | 12 |
| 20 | TFAP2C | T | EZH2 > POLR2A | 81.2 | |
| 6 | TFAP2D | T | EZH2 | 0.4 | 12 |
| 1 | TFAP2E | B | POLR2A + EZH2 | 76.9 | |
| 16 | TFAP4 | B | POLR2A | 18.2 | |
| 12 | TFCP2 | B | POLR2A | 27.6 | |
| 23 | TFCP2L1 | B/T | POLR2A + EZH2 | 146.6 | |
| 13 | TFDP1 | B | POLR2A | 35.0 | |
| 3 | TFDP2 | B | POLR2A | 21.2 | |
| 23 | TFDP3 | T | neither | 9.6 | |
| 23 | TFE3 | B | POLR2A | 96.9 | |
| 6 | TFEB | B | POLR2A | 20.4 | |
| 7 | TFEC | T | neither | 8.6 | |
| 18 | TGIF1 | B | POLR2A | 18.3 | |
| 23 | TGIF2LX | T | POLR2A | 12.6 | |
| 24 | TGIF2LY | T | neither | 5.5 | |
| 15 | THAP10 | B | POLR2A | 9.8 | |
| 1 | THAP3 | B | POLR2A | 18.7 | |
| 17 | THRA | B | POLR2A | 48.6 | |
| 3 | THRB | B | POLR2A + EZH2 | 15.5 | |
| 10 | TLX1 | T | EZH2 | 74.2 | 14 |
| 2 | TLX2 | T | EZH2 | 2.5 | 9 |
| 5 | TLX3 | T | EZH2 | 48.3 | 8 |
| 1 | TOE1 | B | POLR2A | 18.0 | |
| 8 | TOX | T | POLR2A + EZH2 | 22.4 | |
| 20 | TOX2 | B | EZH2 + POLR2A | 25.2 | |
| 16 | TOX3 | T | POLR2A + EZH2 | 11.0 | |
| 14 | TOX4 | B | POLR2A | 44.6 | |
| 17 | TP53 | B | POLR2A | 32.4 | |
| 3 | TP63 | T | POLR2A | 138.9 | |
| 1 | TP73 | T | POLR2A | 11.0 | |
| 19 | TPRX1 | T | neither | 2.8 | |
| 6 | TRERF1 | B | POLR2A | 10.1 | |
| 8 | TRPS1 | B | POLR2A | 15.4 | |
| 13 | TSC22D1 | B | POLR2A | 166.2 | |
| 23 | TSC22D3 | B | POLR2A | 404.8 | |

| | | | | | |
|---|---|---|---|---|---|
| 7 | TSC22D4 | B | POLR2A | 258.4 | |
| 18 | TSHZ1 | B | POLR2A | 37.3 | |
| 20 | TSHZ2 | T | POLR2A | 14.0 | |
| 19 | TSHZ3 | T | POLR2A + EZH2 | 67.3 | |
| 11 | TUB | B | EZH2 | 43.7 | 10 |
| 7 | TWIST1 | T | EZH2 > POLR2A | 42.0 | |
| 3 | UBP1 | B | POLR2A | 54.8 | |
| 17 | UBTF | B | POLR2A | 43.2 | |
| 7 | UNCX | T | EZH2 | 43.4 | 10 |
| 16 | UNKL | B | POLR2A | 17.0 | |
| 1 | USF1 | B | POLR2A | 64.7 | |
| 19 | USF2 | B | POLR2A | 127.8 | |
| 10 | VAX1 | T | EZH2 | 1.6 | 12 |
| 2 | VAX2 | T/B | EZH2 + POLR2A | 19.2 | |
| 12 | VDR | B | POLR2A + EZH2 | 17.9 | |
| 10 | VENTX | B/T | EZH2 | 6.7 | 4 |
| 17 | VEZF1 | B | POLR2A | 42.3 | |
| 20 | VSX1 | T | EZH2 | 20.9 | 8 |
| 14 | VSX2 | T | EZH2 | 0.7 | 19 |
| 14 | WDHD1 | B | POLR2A | 2.5 | |
| 19 | WIZ | B | POLR2A > EZH2 | 32.0 | |
| 11 | WT1 | T | EZH2 | 109.8 | 12 |
| 22 | XBP1 | B | POLR2A | 286.0 | |
| 1 | YBX1 | B | POLR2A | 617.0 | |
| 1 | YBX2 | T | POLR2A | 1335.0 | 5 |
| 12 | YBX3 | B | POLR2A | 250.4 | |
| 14 | YY1 | B | POLR2A | 34.3 | |
| 23 | YY2 | B | neither | 7.3 | |
| 23 | ZBED1 | B | POLR2A | 0.1 | |
| 22 | ZBED4 | B | POLR2A > EZH2 | 8.7 | |
| 20 | ZBP1 | T | EZH2 + POLR2A | 17.6 | |
| 14 | ZBTB1 | B | POLR2A | 31.1 | |
| 3 | ZBTB11 | B | POLR2A | 11.7 | |
| 18 | ZBTB14 | B | POLR2A | 10.8 | |
| 11 | ZBTB16 | B | EZH2 > POLR2A | 35.6 | |
| 1 | ZBTB17 | B | POLR2A | 33.8 | |
| 1 | ZBTB18 | B | POLR2A | 371.5 | |
| 6 | ZBTB2 | B | POLR2A | 14.9 | |
| 3 | ZBTB20 | B | POLR2A | 1.4 | |
| 21 | ZBTB21 | B | POLR2A | 9.1 | |
| 6 | ZBTB24 | B | POLR2A | 7.8 | |
| 19 | ZBTB32 | T | POLR2A | 109.3 | |
| 23 | ZBTB33 | B | POLR2A | 12.3 | |
| 9 | ZBTB34 | B | POLR2A | 11.5 | |
| 3 | ZBTB38 | B | POLR2A | 18.8 | |

| | | | | | |
|---|---|---|---|---|---|
| 12 | ZBTB39 | B | POLR2A | 5.1 | |
| 17 | ZBTB4 | B | POLR2A | 76.3 | |
| 1 | ZBTB41 | B | POLR2A | 9.2 | |
| 9 | ZBTB43 | B | POLR2A | 13.3 | |
| 19 | ZBTB45 | B | POLR2A | 14.9 | |
| 20 | ZBTB46 | B | EZH2 + POLR2A | 16.1 | |
| 3 | ZBTB47 | B | POLR2A | 75.9 | |
| 1 | ZBTB48 | B | POLR2A | 29.7 | |
| 9 | ZBTB5 | B | POLR2A | 19.4 | |
| 19 | ZBTB7A | B | POLR2A | 33.5 | |
| 1 | ZBTB7B | B | POLR2A | 69.5 | |
| 1 | ZC3H11A | B | POLR2A | 2.2 | |
| 2 | ZC3H15 | B | POLR2A | 47.9 | |
| 8 | ZC3H3 | B | POLR2A | 30.2 | |
| 9 | TUT7 | B | POLR2A | 16.1 | |
| 10 | ZEB1 | B | POLR2A | 40.8 | |
| 2 | ZEB2 | B | EZH2 + POLR2A | 26.5 | |
| 8 | ZFAT | B | POLR2A | 8.3 | |
| 14 | ZFHX2 | B | POLR2A | 4.2 | |
| 16 | ZFHX3 | B | EZH2 + POLR2A | 26.0 | |
| 9 | ZFP37 | B | EZH2 > POLR2A | 3.9 | |
| 20 | ZFP64 | B | POLR2A | 6.8 | |
| 16 | ZFPM1 | B | POLR2A | 5.9 | |
| 8 | ZFPM2 | B | POLR2A > EZH2 | 8.6 | |
| 23 | ZFX | B | POLR2A | 12.7 | |
| 24 | ZFY | B | POLR2A | 5.1 | |
| 14 | ZFYVE26 | B | POLR2A | 10.7 | |
| 8 | ZHX1 | B | POLR2A | 17.6 | |
| 8 | ZHX2 | B | EZH2 + POLR2A | 40.5 | |
| 20 | ZHX3 | B | POLR2A | 26.4 | |
| 3 | ZIC1 | T | EZH2 | 426.0 | 8 |
| 13 | ZIC2 | T | POLR2A+ EZH2+ | 298.1 | |
| 23 | ZIC3 | T | EZH2 | 27.6 | 6 |
| 3 | ZIC4 | T | EZH2 | 164.0 | 4 |
| 13 | ZIC5 | T | EZH2 | 53.9 | 6 |
| 19 | ZIK1 | B | POLR2A + EZH2 | 4.6 | |
| 19 | ZIM3 | T | neither | 1.0 | |
| 19 | ZKSCAN1 | B | POLR2A | 35.1 | |
| 16 | ZKSCAN2 | B | POLR2A | 2.8 | |
| 6 | ZKSCAN3 | B | POLR2A | 6.7 | |
| 11 | ZKSCAN4 | B | POLR2A | 8.8 | |
| 7 | ZKSCAN5 | B | POLR2A | 8.5 | |
| 23 | ZMAT1 | B | POLR2A | 35.8 | |
| 3 | ZMAT3 | B | POLR2A | 14.3 | |
| 12 | ZNF10 | B | POLR2A | 12.5 | |

| | | | | | |
|---|---|---|---|---|---|
| 7 | ZNF12 | B | POLR2A | 16.1 | |
| 5 | ZNF131 | B | POLR2A | 12.0 | |
| 19 | ZNF134 | B | POLR2A | 10.1 | |
| 19 | ZNF136 | B | POLR2A | 10.5 | |
| 11 | ZNF143 | B | POLR2A | 13.9 | |
| 19 | ZNF146 | B | POLR2A | 57.0 | |
| 3 | ZNF148 | B | POLR2A | 15.9 | |
| 23 | ZNF157 | T | neither | 2.6 | |
| 8 | ZNF16 | B | POLR2A | 13.3 | |
| 16 | ZNF174 | B | POLR2A | 11.3 | |
| 19 | ZNF175 | B | POLR2A | 8.2 | |
| 6 | ZNF184 | B | POLR2A | 8.0 | |
| 11 | ZNF202 | B | POLR2A | 12.5 | |
| 20 | ZNF217 | B | POLR2A | 20.9 | |
| 14 | ZNF219 | B | POLR2A | 39.2 | |
| 19 | ZNF224 | B | POLR2A | 19.2 | |
| 18 | ZNF24 | B | POLR2A | 26.2 | |
| 8 | ZNF251 | B | POLR2A | 48.1 | |
| 19 | ZNF256 | B | POLR2A | 8.3 | |
| 19 | ZNF260 | B | POLR2A | 7.8 | |
| 16 | ZNF276 | B | POLR2A | 27.4 | |
| 22 | ZNF280A | T | neither | 1.9 | |
| 1 | ZNF281 | B | POLR2A | 11.5 | |
| 7 | ZNF282 | B | POLR2A | 29.8 | |
| 6 | ZNF292 | B | POLR2A | 14.2 | |
| 19 | ZNF296 | B | POLR2A | 9.2 | |
| 5 | ZNF300 | B | POLR2A | 8.9 | |
| 19 | ZNF304 | B | POLR2A | 9.3 | |
| 16 | ZNF319 | B | POLR2A | 12.1 | |
| 10 | ZNF32 | B | POLR2A | 86.5 | |
| 6 | ZNF322 | B | POLR2A | 4.6 | |
| 19 | ZNF333 | B | POLR2A | 15.2 | |
| 20 | ZNF335 | B | POLR2A | 21.8 | |
| 20 | ZNF341 | B | POLR2A | 6.8 | |
| 19 | ZNF350 | B | POLR2A | 9.5 | |
| 5 | ZNF354A | B | POLR2A | 12.4 | |
| 5 | ZNF354B | B | POLR2A | 15.7 | |
| 5 | ZNF354C | B | POLR2A | 6.3 | |
| 1 | ZNF362 | B | POLR2A | 46.3 | |
| 10 | ZNF365 | T | EZH2 | 36.5 | 10 |
| 5 | ZNF366 | B | POLR2A | 2.4 | |
| 9 | ZNF367 | B | POLR2A | 3.2 | |
| 10 | ZNF37A | B | POLR2A | 9.5 | |
| 19 | ZNF382 | B | POLR2A | 5.7 | |
| 12 | ZNF384 | B | POLR2A | 43.6 | |

| | | | | | |
|---|---|---|---|---|---|
| 7 | ZNF394 | B | POLR2A | 23.5 | |
| 7 | ZNF398 | B | POLR2A | 11.0 | |
| 14 | ZNF410 | B | POLR2A | 17.9 | |
| 19 | ZNF415 | B | POLR2A | 9.0 | |
| 19 | ZNF418 | B | POLR2A | 6.0 | |
| 16 | ZNF423 | B | POLR2A | 4.9 | |
| 7 | ZNF425 | B | POLR2A | 3.8 | |
| 19 | ZNF426 | B | POLR2A | 5.8 | |
| 19 | ZNF430 | B | POLR2A | 4.2 | |
| 19 | ZNF431 | B | POLR2A | 7.2 | |
| 10 | ZNF438 | B | POLR2A | 14.9 | |
| 19 | ZNF444 | B | POLR2A | 38.7 | |
| 3 | ZNF445 | B | POLR2A | 5.5 | |
| 7 | ZNF467 | B | POLR2A | 21.0 | |
| 19 | ZNF480 | B | POLR2A | 6.0 | |
| 10 | ZNF488 | T | POLR2A | 83.6 | |
| 1 | ZNF496 | B | POLR2A | 17.9 | |
| 4 | ZBTB49 | B | POLR2A | 5.9 | |
| 9 | ZNF510 | B | POLR2A | 7.4 | |
| 2 | ZNF513 | B | POLR2A | 46.8 | |
| 18 | ZNF516 | B | POLR2A | 30.0 | |
| 18 | ZNF521 | B | EZH2 | 12.3 | 10 |
| 19 | ZNF536 | T | POLR2A | 17.1 | |
| 19 | ZNF551 | B | POLR2A | 9.7 | |
| 19 | ZNF568 | B | POLR2A | 3.5 | |
| 1 | ZNF593 | B | POLR2A | 1.3 | |
| 16 | ZNF598 | B | POLR2A | 54.0 | |
| 19 | ZNF606 | B | POLR2A | 7.8 | |
| 5 | ZNF608 | B | POLR2A | 11.5 | |
| 15 | ZNF609 | B | POLR2A | 19.8 | |
| 5 | ZNF622 | B | POLR2A | 34.0 | |
| 19 | ZNF628 | B | POLR2A | 8.2 | |
| 2 | ZNF638 | B | POLR2A | 47.5 | |
| 12 | ZNF641 | B | POLR2A | 13.7 | |
| 1 | ZPF69 | B | POLR2A | 3.9 | |
| 1 | ZNF644 | B | POLR2A | 13.8 | |
| 19 | ZNF649 | B | POLR2A | 6.0 | |
| 17 | ZNF652 | B | POLR2A | 15.7 | |
| 19 | ZNF653 | B | POLR2A | 9.7 | |
| 9 | ZNF658 | B | POLR2A | 2.1 | |
| 1 | ZNF683 | T | neither | 126.1 | |
| 1 | ZNF691 | B | POLR2A | 17.4 | |
| 19 | ZNF699 | B | POLR2A | 2.9 | |
| 8 | ZNF703 | B | POLR2A | 73.9 | |
| 8 | ZNF704 | B | POLR2A | 12.9 | |

| | | | | |
|---|---|---|---|---|
| 8 | ZNF706 | B | POLR2A | 44.0 |
| 15 | ZNF710 | B | POLR2A | 15.7 |
| 23 | ZNF711 | B | POLR2A | 19.4 |
| 12 | ZNF740 | B | POLR2A | 21.3 |
| 7 | ZNF746 | B | POLR2A | 9.6 |
| 17 | ZNF750 | T | POLR2A | 204.1 |
| 7 | ZNF777 | B | POLR2A | 19.6 |
| 9 | ZNF782 | B | POLR2A | 3.6 |
| 7 | ZNF786 | B | POLR2A | 8.3 |
| 19 | ZNF8 | B | POLR2A | 5.5 |
| 16 | ZNF821 | B | POLR2A | 11.6 |
| 19 | ZNF829 | B | POLR2A | 2.2 |
| 17 | ZNF830 | B | POLR2A | 20.8 |
| 20 | ZNF831 | T | POLR2A | 4.2 |
| 19 | ZNF837 | B | POLR2A | 5.6 |
| 19 | ZNF845 | B | POLR2A | 4.4 |
| 19 | ZNF91 | B | POLR2A | 7.6 |
| 19 | ZNF93 | B | POLR2A | 2.9 |
| 19 | ZSCAN1 | T | POLR2A | 11.6 |
| 16 | ZSCAN10 | T | POLR2A | 0.5 |
| 6 | ZSCAN12 | B | neither | 7.4 |
| 6 | ZSCAN16 | B | POLR2A | 11.5 |
| 15 | ZSCAN2 | B | POLR2A | 4.6 |
| 19 | ZSCAN20 | B | POLR2A | 2.3 |
| 7 | ZSCAN21 | B | POLR2A | 11.6 |
| 19 | ZSCAN22 | B | POLR2A | 3.6 |
| 6 | ZSCAN23 | B | POLR2A | 1.7 |
| 15 | ZSCAN29 | B | POLR2A | 12.1 |
| 19 | ZSCAN4 | T | neither | 0.7 |
| 19 | ZSCAN5A | T | POLR2A | 2.2 |
| 23 | ZXDA | B | POLR2A | 3.8 |
| 23 | ZXDB | B | POLR2A | 6.4 |
| 1 | ZZZ3 | B | POLR2A | 14.6 |

B: broadly expressed gene; T tissue targeted gene; Gene Expression: transcripts per million (TPM); EZH2 binding in ChIP experiment; signal p-value
