## Supplementary Table2 for "Role of polycomb repressive complex 2 in regulation of human transcription factor gene expression"

Supplementary Table 2: Control; sample of all human genes.
Sequential genes beginning with chr9 ACO1 (183 genes)

| Index | Gene | B/T | POLR2A/EZH2 |
|---|---|---|---|
| 185 | ABCA1 | B | POLR2A |
| 86 | ABHD17B | B | POLR2A |
| 1 | ACO1 | B | POLR2A |
| 115 | AGTPBP1 | B | POLR2A |
| 90 | ALDH1A1 | B | POLR2A |
| 72 | ALDH1B1 | B | POLR2A |
| 75 | ANKRD18A | T | POLR2A + EZH2 |
| 14 | ANKRD18B | T | EZH2 + POLR2A |
| 170 | ANKS6 | B | POLR2A |
| 165 | ANP32B | B | POLR2A |
| 91 | ANXA1 | B | POLR2A |
| 147 | AOPEP | B | POLR2A |
| 4 | APTX | B | POLR2A |
| 11 | AQP7 | T | POLR2A |
| 26 | ARID3C | T | EZH2 |
| 126 | AUH | B | POLR2A |
| 7 | B4GALT1 | B | POLR2A |
| 181 | BAAT | T | POLR2A |
| 8 | BAG1 | B | POLR2A |
| 134 | BICD2 | B | POLR2A |
| 53 | C9orf128 | B | POLR2A |
| 33 | C9orf131 | T | neither |
| 111 | C9orf64 | B | POLR2A |
| 87 | C9orf85 | B | POLR2A |
| 44 | CA9 | T | POLR2A |
| 94 | CARNMT1 | B | POLR2A |
| 179 | CAVIN4 | T | POLR2A |
| 43 | CCDC107 | B | POLR2A |
| 158 | CCDC180 | B | POLR2A |
| 41 | CD72 | T | POLR2A |
| 155 | CDC14B | B | POLR2A- |
| 131 | CENPP | B | POLR2A |
| 9 | CHMP5 | B | POLR2A |
| 57 | CLTA | B | POLR2A |
| 24 | CNTFR | B | EZH2 |
| 77 | CNTNAP3 | B | neither |
| 172 | COL15A1 | B | EZH2 |
| 167 | CORO2A | T | POLR2A |
| 47 | CREB3 | B | POLR2A |
| 119 | DAPK1 | B | POLR2A |
| 69 | DCAF10 | B | POLR2A |

| | | | |
|---|---|---|---|
| 18 | DCAF12 | B | POLR2A |
| 25 | DCTN3 | B | POLR2A |
| 2 | DDX58 | B | POLR2A |
| 23 | DNAI1 | T | POLR2A |
| 5 | DNAJA1 | B | POLR2A |
| 32 | DNAJB5 | B | POLR2A |
| 132 | ECM2 | B | neither |
| 150 | ERCC6L2 | B | POLR2A |
| 176 | ERP44 | B | POLR2A |
| 140 | FAM120A | B | POLR2A |
| 29 | FAM205A | T | EZH2 |
| 30 | FAM205C | T | neither |
| 36 | FAM214B | B | POLR2A |
| 22 | FAM219A | B | POLR2A |
| 54 | FAM221B | T | POLR2A |
| 76 | FAM240B | T | neither |
| 74 | FAM95C | B | POLR2A |
| 148 | FANCC | B | POLR2A |
| 146 | FBP1 | T | EZH2 + POLR2A |
| 66 | FBXO10 | B | POLR2A |
| 136 | FGD3 | B | POLR2A |
| 106 | FRMD3 | B | EZH2 + POLR2A |
| 67 | FRMPD1 | T | EZH2 |
| 187 | FSD1L | B | POLR2A |
| 169 | GABBR2 | T | EZH2 |
| 171 | GALNT12 | T | POLR2A |
| 27 | GALT | B | POLR2A |
| 48 | GBA2 | B | POLR2A |
| 98 | GCNT1 | B | POLR2A |
| 88 | GDA | T | POLR2A + EZH2 |
| 109 | GKAP1 | B | POLR2A |
| 56 | GLIPR2 | B | EZH2 + POLR2A |
| 101 | GNA14 | B | EZH2 |
| 102 | GNAQ | B | POLR2A |
| 58 | GNE | B | POLR2A |
| 117 | GOLM1 | B | POLR2A + EZH2 |
| 63 | GRHPR | B | POLR2A |
| 184 | GRIN3A | T | EZH2 |
| 154 | HABP4 | B | POLR2A |
| 151 | HSD17B3 | B | neither |
| 129 | IARS1 | B | POLR2A |
| 107 | IDNK | B | POLR2A |
| 73 | IGFBPL1 | T | EZH2 |
| 28 | IL11RA | B | POLR2A |
| 177 | INVS | B | POLR2A |

| | | | |
|---|---|---|---|
| 133 | IPPK | B | POLR2A |
| 80 | KGFLP2 | B | neither |
| 31 | KIAA1045 | T | EZH2 |
| 20 | KIF24 | B | POLR2A |
| 110 | KIF27 | B | POLR2A |
| 83 | KLF9 | B | POLR2A |
| 81 | MAMDC2 | T | POLR2A |
| 60 | MELK | T | POLR2A |
| 144 | MFSD14B | B | POLR2A |
| 116 | NAA35 | B | POLR2A |
| 161 | NCBP1 | B | POLR2A |
| 3 | NDUFB6 | B | POLR2A |
| 10 | NFX | B | POLR2A |
| 138 | NINJ1 | B | POLR2A |
| 95 | NMRK1 | B | POLR2A |
| 12 | NOL6 | B | POLR2A |
| 130 | NOL8 | B | POLR2A |
| 50 | NPR2 | B | POLR2A- |
| 174 | NR4A3 | T | EZH2 +POLR2A |
| 114 | NTRK2 | B | EZH2 |
| 21 | NUDT2 | B | POLR2A |
| 143 | NUTM2F | T | neither |
| 96 | OSTF1 | B | POLR2A |
| 61 | PAX5 | T | EZH2 |
| 145 | PCAT7 | T | POLR2A |
| 97 | PCSK5 | B | POLR2A |
| 141 | PHF2 | B | POLR2A |
| 35 | PIGO | B | POLR2A |
| 180 | PLPPR1 | T | EZH2 |
| 65 | POLR1E | B | POLR2A |
| 15 | PRSS3 | T | POLR2A |
| 99 | PRUNE2 | B | POLR2A |
| 149 | PTCH1 | B | POLR2A |
| 163 | PTCSC2 | T | EZH2 |
| 142 | PTPDC1 | B | POLR2A |
| 105 | RASEF | T | EZH2 + POLR2A |
| 55 | RECK | B | POLR2A |
| 49 | RGP1 | B | POLR2A |
| 112 | RMI1 | B | POLR2A |
| 183 | RNF20 | B | POLR2A |
| 59 | RNF38 | B | POLR2A |
| 127 | ROR2 | T | EZH2 + POLR2A |
| 92 | RORB | T | EZH2 |
| 39 | RUSC2 | B | POLR2A |
| 123 | SEMA4D | B | POLR2A |

| | | | |
|---|---|---|---|
| 71 | SHB | B | EZH2 + POLR2A |
| 122 | SHC3 | T | EZH2 + POLR2A |
| 42 | SIT1 | T | POLR2A |
| 70 | SLC25A51 | B | POLR2A |
| 113 | SLC28A3 | T | POLR2A |
| 152 | SLC35D2 | B | POLR2A |
| 186 | SLC44A1 | B | POLR2A |
| 82 | SMC5 | B | POLR2A |
| 6 | SMU1 | B | POLR2A |
| 51 | SPAG8 | B | POLR2A |
| 121 | SPIN1 | B | POLR2A |
| 128 | SPTLC1 | B | POLR2A |
| 175 | STX17 | B | POLR2A |
| 13 | SUGT1P1 | B | POLR2A |
| 137 | SUSD3 | B | POLR2A |
| 125 | SYK | B | EZH2 |
| 168 | TBC1D2 | B | POLR2A |
| 159 | TDRD7 | B | POLR2A |
| 40 | TESK1 | B | POLR2A |
| 178 | TEX10 | B | POLR2A |
| 173 | TGFBR1 | B | POLR2A |
| 104 | TLE1 | B | POLR2A |
| 103 | TLE4 | B | POLR2A |
| 46 | TLN1 | B | POLR2A |
| 89 | TMC1 | T | POLR2A |
| 85 | TMEM2 | B | POLR2A |
| 52 | TMEM8B | B | POLR2A |
| 45 | TPM2 | B | POLR2A |
| 166 | TRIM14 | B | POLR2A |
| 164 | TRMO | B | POLR2A |
| 68 | TRMT10B | B | POLR2A |
| 84 | TRPM3 | T | EZH2 |
| 93 | TRPM6 | B | POLR2A |
| 160 | TSTD2 | B | POLR2A |
| 118 | TUT7 | B | POLR2A |
| 19 | UBAP1 | B | POLR2A |
| 17 | UBAP2 | B | POLR2A |
| 16 | UBE2R2 | B | POLR2A |
| 108 | UBQLN1 | B | POLR2A |
| 37 | UNC13B | B | POLR2A |
| 124 | UNQ6494 | T | POLR2A |
| 34 | VCP | B | POLR2A |
| 100 | VPS13A | B | POLR2A + EZH2 |
| 139 | WNK2 | T | POLR2A + EZH2 |
| 162 | XPA | B | POLR2A |

| | | | |
|---|---|---|---|
| 64 | ZBTB5 | B | POLR2A |
| 62 | ZCCHC7 | B | POLR2A |
| 182 | ZNF189 | B | POLR2A |
| 153 | ZNF367 | B | POLR2A |
| 135 | ZNF484 | B | POLR2A |
| 78 | ZNF658 | B | POLR2A |
| 156 | ZNF782 | B | POL |
