## Supplementary Table3 for "Role of polycomb repressive complex 2 in regulation of human transcription factor gene expression"

Supplementary Table 3: All database testis-specific genes with information about prominent transcription factors in the promoter. (204 genes)

| Index | Gene | Chr | Prominent TF in Promoter |
|---|---|---|---|
| 127 | ACRV1 | 11 | FOS |
| 111 | ACTL7A | 9 | ATF7 |
| 105 | ACTL7B | 9 | CREB1 |
| 213 | ACTL9 | 19 | CREB1 |
| 16 | ACTRT2 | 1 | EZH2 |
| 55 | ADAD1 | 4 | HMBOX1 |
| 101 | ADAM2 | 8 | RAD21 |
| 64 | ADAM29 | 4 | IKZF1 |
| 4 | ADAM30 | 1 | ZBTB33 |
| 21 | ADCY10 | 1 | YY1 |
| 239 | AKAP4 | 23 | GATA1 |
| 42 | ALS2CR12 | 2 | ASH2L |
| 26 | APOBEC4 | 1 | IKZF1 |
| 118 | ASTN2-AS1 | 9 | ATF2 |
| 28 | ATP1A4 | 1 | FOXA1 |
| 10 | AXDND1 | 1 | POLR2A |
| 203 | BOD1L2 | 18 | ZBTB33 |
| 33 | BOLL | 2 | EZH2 |
| 7 | BRDT | 1 | ZBTB33 |
| 176 | C16orf82 | 16 | KDM1A |
| 194 | C17orf47 | 17 | EZH2 |
| 5 | C1orf94 | 1 | POLR2A |
| 34 | C2CD6 | 2 | ZBTB33 |
| 38 | C2orf78 | 2 | none |
| 66 | C5orf47 | 5 | PCBP1 |
| 70 | C5orf60 | 5 | MNT |
| 85 | C6orf10 | 6 | MAFK |
| 108 | C9orf131 | 9 | ATF7 |
| 107 | C9orf135-AS1 | 9 | ETS1 |
| 65 | CABS1 | 4 | CEBPB |
| 199 | CABYR | 18 | PHF8 |
| 206 | CALR3 | 19 | POLR2A |
| 179 | CASC16 | 16 | POLR2A |
| 14 | CATSPERE | 1 | POLR2A |
| 59 | CCDC110 | 4 | PHF8 |
| 204 | CCDC114 | 19 | POLR2A |
| 228 | CCDC116 | 22 | TBL1XR1 |
| 13 | CCDC185 | 1 | EZH2 |
| 20 | CCDC27 | 1 | none |
| 43 | CCDC36 | 3 | POLR2A |

| | | | |
|---|---|---|---|
| 48 | CCDC37-AS1 | 3 | ASH2L |
| 134 | CCDC38 | 12 | POLR2A |
| 186 | CCDC42 | 17 | POLR2A |
| 133 | CCDC62 | 12 | POLR2A |
| 132 | CCDC63 | 12 | EZH2 |
| 120 | CCDC7 | 10 | POLR2A |
| 172 | TERB1 | 16 | POLR2A |
| 125 | CCDC83 | 11 | POLR2A |
| 135 | CCER1 | 12 | ZBTB33 |
| 114 | CCIN | 9 | ATF2 |
| 138 | CEP83-AS1 | 12 | POLR2A |
| 200 | CETN1 | 18 | XRCC5 |
| 80 | CFAP206 | 6 | KDM4A |
| 201 | CFAP53 | 18 | POLR2A |
| 90 | CPA5 | 7 | POLR2A |
| 54 | CPEB2-AS1 | 4 | POLR2A |
| 86 | CRISP2 | 6 | ZBTB33 |
| 160 | CT62 | 15 | POLR2A |
| 234 | CYLC1 | 23 | ATF2 |
| 35 | DAW1 | 2 | POLR2A |
| 45 | DAZL | 3 | NFATC1 |
| 53 | DCAF4L1 | 4 | SUZ12 |
| 96 | DCAF4L2 | 8 | NRF1 |
| 67 | DDX4 | 5 | RELB |
| 106 | DMRT1 | 9 | EZH2 |
| 3 | DMRTB1 | 1 | NRF1 |
| 211 | DMRTC2 | 19 | ZBTB33 |
| 193 | DNAH9 | 17 | POLR2A |
| 102 | DNAJC5B | 8 | POLR2A |
| 171 | DPEP3 | 16 | CTCF |
| 75 | EGFLAM-AS2 | 5 | FOS |
| 223 | EPPIN | 20 | SMC3 |
| 159 | EXD1 | 15 | POLR2A |
| 51 | F11-AS1 | 4 | POLR2A |
| 144 | FAM186B | 12 | ATF2 |
| 214 | FAM187B | 19 | POLR2A |
| 110 | FAM205A | 9 | none |
| 77 | FAM217A | 6 | FOS |
| 232 | FAM46D | 23 | POLR2A |
| 230 | FAM47B | 23 | none |
| 231 | FAM47C | 23 | none |
| 24 | FAM71A | 1 | none |
| 74 | FAM71B | 5 | ATF2 |
| 154 | FAM71D | 14 | FOS |
| 89 | FBXO24 | 7 | ZBTB33 |

| | | | |
|---|---|---|---|
| 198 | FBXW10 | 17 | CHD7 |
| 88 | FKBP6 | 7 | none |
| 153 | FSCB | 14 | CREB1 |
| 32 | FSIP2 | 2 | TRIM28 |
| 94 | GALNTL5 | 7 | TRIM28 |
| 205 | GGN | 19 | POLR2A |
| 60 | GK2 | 4 | ZBTB33 |
| 164 | GOLGA8F | 15 | none |
| 167 | GOLGA8G | 15 | none |
| 163 | GOLGA8S | 15 | CTCF |
| 136 | GSG1 | 12 | POLR2A |
| 130 | H1FNT | 12 | none |
| 76 | HDGFL1 | 6 | CTCF |
| 157 | HEATR4 | 14 | POLR2A |
| 197 | HEATR9 | 17 | none |
| 17 | HORMAD1 | 1 | ZBTB33 |
| 225 | HORMAD2 | 22 | ATF7 |
| 99 | KCNU1 | 8 | PCBP2 |
| 196 | KIF2B | 17 | CREB1 |
| 22 | KLF17 | 1 | ZBTB33 |
| 137 | KRT72 | 12 | ZBTB33 |
| 158 | LDHAL6B | 15 | NRF1 |
| 126 | LDHC | 11 | POLR2A |
| 9 | LRRC71 | 1 | EZH2 |
| 156 | LRRC74A | 14 | EHMT2 |
| 236 | MAGEB3 | 23 | REST |
| 237 | MAGEB4 | 23 | EBF1 |
| 238 | MAGEC2 | 23 | POLR2A |
| 192 | MARCH10 | 17 | EZH2 |
| 103 | MCMDC2 | 8 | POLR2A |
| 185 | MEIOC | 17 | EZH2 |
| 46 | MORC1 | 3 | CTCF |
| 73 | MROH2B | 5 | FOS |
| 104 | MROH5 | 8 | POLR2A |
| 188 | MYCBPAP | 17 | KDM4A |
| 25 | NBPF4 | 1 | none |
| 216 | NLRP11 | 19 | POLR2A |
| 209 | NLRP4 | 19 | PKNOX1 |
| 91 | NME8 | 7 | RAD21 |
| 23 | NUP210L | 1 | POLR2A |
| 166 | NUTM1 | 15 | IKZF1 |
| 81 | OR2H1 | 6 | EGR1 |
| 165 | OR4N4 | 15 | CTCF |
| 180 | OTOA | 16 | IKZF1 |
| 143 | OVOS2 | 12 | ASH2L |

| | | | |
|---|---|---|---|
| 1 | OXCT2 | 1 | EZH2 |
| 87 | PAPOLB | 7 | POLR2A |
| 235 | PASD1 | 23 | MNT |
| 63 | PDHA2 | 4 | ATF2 |
| 83 | PGK2 | 6 | PKNOX1 |
| 141 | PIWIL1 | 12 | CTCF |
| 142 | PLCZ1 | 12 | FOS |
| 183 | PMFBP1 | 16 | IKZF1 |
| 113 | PPP3R2 | 9 | REST |
| 71 | PRDM9 | 5 | RAD21 |
| 184 | PRM1 | 16 | SRF |
| 170 | PRM2 | 16 | AGO1 |
| 222 | PRND | 20 | FOS |
| 41 | PRR30 | 2 | none |
| 182 | PRSS54 | 16 | ATF2 |
| 58 | RBM46 | 4 | SMC3 |
| 229 | RBMXL3 | 23 | MAFK |
| 227 | RFPL3S | 22 | ZBTB33 |
| 140 | RFX4 | 12 | EZH2 |
| 18 | RGSL1 | 1 | CTCF |
| 226 | RIMBP3 | 22 | none |
| 146 | RNF17 | 13 | EZH2 |
| 47 | ROPN1B | 3 | EZH2 |
| 155 | RPGRIP1 | 14 | EZH2 |
| 212 | RSPH6A | 19 | POLR2A |
| 177 | SEPT12 | 16 | ZBTB33 |
| 95 | SEPT14 | 7 | ATF2 |
| 37 | SH2D6 | 2 | POLR2A |
| 12 | SHCBP1L | 1 | ZBTB7B |
| 218 | SIGLECL1 | 19 | SMC3 |
| 56 | SLC25A31 | 4 | ZBTB33 |
| 69 | SLC36A3 | 5 | IKZF1 |
| 217 | SLC6A16 | 19 | TRIM22 |
| 19 | SLC9C2 | 1 | POLR2A |
| 68 | SLCO6A1 | 5 | CEBPB |
| 27 | SMCP | 1 | CTCF |
| 119 | SPAG6 | 10 | EZH2 |
| 92 | SPAM1 | 7 | FOS |
| 49 | SPATA16 | 3 | POLR2A |
| 189 | SPATA22 | 17 | SP1 |
| 117 | SPATA31E1 | 9 | CTCF |
| 169 | SPATA8 | 15 | ZFX |
| 100 | SPATC1 | 8 | POLR2A |
| 149 | SPERT | 13 | TRIM22 |
| 2 | SYCP1 | 1 | EZH2 |

| | | | |
|---|---|---|---|
| 78 | TCP11 | 6 | ASH2L |
| 79 | TCTE1 | 6 | EZH2 |
| 123 | TDRD1 | 10 | MNT |
| 11 | TDRD5 | 1 | KDM1A |
| 187 | TEKT3 | 17 | EZH2 |
| 178 | TEKT5 | 16 | POLR2A |
| 190 | TEX14 | 17 | POLR2A |
| 52 | TKTL2 | 4 | MGA |
| 168 | TMCO5A | 15 | MAFK |
| 131 | TMPRSS12 | 12 | ASH2L |
| 109 | TPD52L3 | 9 | REST |
| 224 | TPTE | 21 | none |
| 44 | TRIM42 | 3 | CHD4 |
| 62 | TRIML1 | 4 | REST |
| 8 | TSACC | 1 | POLR2A |
| 129 | TSGA10IP | 11 | RBBP5 |
| 36 | TSPYL6 | 2 | POLR2A |
| 72 | TSSK1B | 5 | none |
| 57 | TTC29 | 4 | SIN3A |
| 15 | TTLL10 | 1 | ZBTB33 |
| 84 | TTLL2 | 6 | EZH2 |
| 148 | TUBA3C | 13 | ZBTB33 |
| 29 | TUBA3E | 2 | ZBTB33 |
| 50 | TUSC7 | 3 | CREB1 |
| 202 | TXNDC2 | 18 | POLR2A |
| 128 | UBQLN3 | 11 | CREB1 |
| 139 | USP44 | 12 | POLR2A |
| 208 | WDR87 | 19 | ASH2L |
| 191 | ZPBP2 | 17 | IKZF1 |
