## Supplementary Table4 for "Role of polycomb repressive complex 2 in regulation of human transcription factor gene expression"

Supplementary Table 4: All database human transcription factor genes with annotated protein structural elements (1020 genes)

| Chr | Gene | B/T | POLR2A/EZH2 | Structure |
|---|---|---|---|---|
| 20 | ADNP | B | POLR2A | homeobox |
| 18 | ADNP2 | B | POLR2A | homeobox |
| 2 | AFF3 | T | EZH2 | |
| 5 | AFF4 | B | POLR2A | P-TEFB complex |
| 12 | ALX1 | T | EZH2 + POLR2A | Homeobox TAAT |
| 1 | ALX3 | B | EZH2 > POLR2A | homeobox |
| 11 | ALX4 | T | EZH2 | homeobox |
| 19 | ARID3A | B/T | neither | ARID family |
| 10 | ARID5B | B | POLR2A | ARID family; histone demethy |
| 1 | ARNT | B | POLR2A | bHLH |
| 15 | ARNT2 | B | EZH2 + POLR2A | bHLH |
| 11 | ARNTL | B | POLR2A > EZH2 | bHLH |
| 12 | ARNTL2 | B | POLR2A | bHLH |
| 12 | ASCL1 | T | EZH2 | bHLH |
| 11 | ASCL2 | T | EZH2 | bHLH |
| 11 | ASCL3 | T | neither | bHLH |
| 12 | ASCL4 | T | EZH2 | bHLH |
| 12 | ATF1 | B | POLR2A | bZIP |
| 2 | ATF2 | B | POLR2A | bZIP |
| 1 | ATF3 | B | POLR2A + EZH2 | CREB family |
| 22 | ATF4 | B | POLR2A | CREB family |
| 19 | ATF5 | B | POLR2A | CRE element |
| 1 | ATF6 | B | POLR2A | cAMP-dependent |
| 6 | ATF6B | B | POLR2A | cAMP-dependent |
| 12 | ATF7 | B | POLR2A | cAMP-dependent |
| 4 | ATOH1 | T | EZH2 | bHLH |
| 10 | ATOH7 | T | POLR2A + EZH2 | bHLH |
| 21 | BACH1 | B | POLR2A | CNC-bZIP |
| 6 | BACH2 | B | POLR2A | CNC-bZIP |
| 9 | BARHL1 | T | EZH2 | homeobox |
| 1 | BARHL2 | T | EZH2 | homeobox |
| 9 | BARX1 | T | EZH2 | homeobox |
| 11 | BARX2 | T | EZH2 | homeobox |
| 14 | BATF | B | EZH2 > POLR2A | bZIP |
| 11 | BATF2 | B/T | POLR2A | bZIP |
| 1 | BATF3 | T | EZH2 | bZIP |
| 2 | BCL11A | T | POLR2A > EZH2 | ZNF |
| 14 | BCL11B | T | POLR2A + EZH2 | ZNF |
| 3 | BCL6 | B | POLR2A | ZNF |
| 17 | BCL6B | B | POLR2A > EZH2 | ZNF |
| 6 | BCLAF1 | B | POLR2A | BCL2-assoc TF |

| | | | | |
|---|---|---|---|---|
| 20 | BHLH23 | T | EZH2 | |
| 3 | BHLHB2 | B | POLR2A | bHLH |
| 7 | BHLHB8 | T | POLR2A | bHLH |
| 8 | BHLHE22 | T | EZH2 | bHLH |
| 15 | BNC1 | T | EZH2 + POLR2A | ZNF |
| 9 | BNC2 | T | POLR2A + EZH2 | ZNF |
| 1 | CAMTA1 | B | POLR2A | Calmodulin binding |
| 17 | CAMTA2 | B | POLR2A | Calmodulin binding |
| 5 | CDX | T | EZH2 | |
| 5 | CDX1 | T | EZH2 | Homeobox |
| 13 | CDX2 | T | EZH2 | Homeobox |
| 19 | CEBPA | B | POLR2A + EZH2 | bZIP recog CCAAT |
| 20 | CEBPB | B | POLR2A | bZIP |
| 19 | CEBPG | B | POLR2A | bZIP |
| 2 | CEBPZ | B | POLR2A | bZIP |
| 9 | CREB3 | B | POLR2A | cAMP resp. bZIP |
| 11 | CREB3L1 | B | POLR2A | cAMP resp. box B |
| 7 | CREB3L2 | B | POLR2A | cAMP resp. bZIP |
| 19 | CREB3L3 | T | neither | cAMP resp. bZIP |
| 1 | CREB3L4 | B | POLR2A | cAMP resp. |
| 7 | CREB5 | B | neither | cAMP resp. bZIP |
| 12 | CREBL2 | B | POLR2A | cAMP resp. bZIP |
| 11 | CREBZF | B | POLR2A | CREB/ATF bZIP |
| 16 | CTCF | B | POLR2A | CCCTC binding ZNF |
| 20 | CTCFL | T | POLR2A | CCCTC binding ZNF |
| 7 | CUX1 | B | POLR2A | Homeobox + CUT |
| 12 | CUX2 | T | POLR2A + EZH2 | Homeobox + CUT |
| 13 | DACH1 | B | neither | Daschund domain |
| 23 | DACH2 | T | neither | Daschund domain |
| 11 | DBX1 | T | EZH2 | Homeobox |
| 12 | DBX2 | T | EZH2 | Homeobox |
| 2 | DLX1 | T | EZH2 | Homeobox |
| 2 | DLX2 | T | EZH2 + POLR2A | Homeobox |
| 17 | DLX3 | T | EZH2 | Homeobox |
| 17 | DLX4 | T | EZH2 | Homeobox |
| 7 | DLX5 | T | EZH2 | Homeobox |
| 7 | DLX6 | T | EZH2 | Homeobox |
| 9 | DMRT1 | T | EZH2- | ZNF + DM |
| 9 | DMRT2 | T | EZH2 | DM |
| 9 | DMRT3 | T | EZH2 | |
| 9 | DMRTA1 | T | POLR2A | |
| 1 | DMRTA2 | T | EZH2 | |
| 1 | DMRTB1 | T | neither | |
| 19 | DMRTC2 | T | neither | |
| 19 | DPF1 | T | POLR2A > EZH2 | Double PHD fingers |

| | | | | |
|---|---|---|---|---|
| 11 | DPF2 | B | POLR2A | ZNF-like |
| 14 | DPF3 | T | EZH2 | Chromatin remodeling |
| 1 | DR1 | B | POLR2A | TBP |
| 11 | DRAP1 | B | POLR2A | Interacts with DR1 |
| 10 | DRGX | T | EZH2 | Homeobox |
| 20 | E2F1 | B | POLR2A | E2F family |
| 1 | E2F2 | T | POLR2A | E2F family |
| 6 | E2F3 | B | POLR2A | E2F family |
| 16 | E2F4 | B | POLR2A | E2F family |
| 8 | E2F5 | B | POLR2A | E2F family |
| 2 | E2F6 | B | POLR2A | E2F family |
| 12 | E2F7 | T | POLR2A | E2F family |
| 11 | E2F8 | T | POLR2A + EZH2 | E2F family |
| 5 | EBF1 | T | EZH2 | ZNF |
| 10 | EBF3 | T | EZH2 | Like EBF1 |
| 20 | EBF4 | B | EZH2 | bHLH |
| 5 | EGR1 | B | POLR2A | ZNF |
| 10 | EGR2 | B | POLR2A > EZH2 | ZNF |
| 2 | EGR4 | T | EZH2 | Similar to EGR2 (ZNF) |
| 13 | ELF1 | B | POLR2A | ETS TF |
| 4 | ELF2 | B | POLR2A | ETS TF |
| 1 | ELF3 | T | POLR2A | ETS TF |
| 23 | ELF4 | B | POLR2A | ETS TF |
| 11 | ELF5 | T | POLR2A | ETS TF |
| 23 | ELK1 | B | POLR2A | ETS TF |
| 12 | ELK3 | B | POLR2A | ETS TF |
| 1 | ELK4 | B | POLR2A | ETS TF |
| 2 | EMX1 | T | EZH2 | Homeobox |
| 10 | EMX2 | T | EZH2 | Homeobox |
| 2 | EN1 | T | EZH2 | Homeobox |
| 7 | EN2 | T | EZH2 | Homeobox |
| 19 | ERF | B | POLR2A | ETS family |
| 21 | ERG | B | EZH2 > POLR2A | ETS family |
| 6 | ESR1 | T | EZH2+POLR2A | Estrogen receptor |
| 14 | ESR2 | T | POLR2A + EZH2 | Estrogen receptor |
| 11 | ESRRA | B | POLR2A | Estrogen receptor related A |
| 14 | ESRRB | T | EZH2 | Estrogen receptor related B |
| 1 | ESRRG | T | EZH2 | Estrogen receptor related G |
| 11 | ETS1 | B | POLR2A | ETS family (GGA(A/T) |
| 21 | ETS2 | B | POLR2A | ETS family |
| 7 | ETV1 | B | POLR2A | ETS family variant |
| 19 | ETV2 | T | POLR2A | ETS family variant |
| 1 | ETV3 | B | POLR2A | ETS family variant |
| 1 | ETV3L | T | neither | ETS family variant |
| 17 | ETV4 | B | POLR2A | ETS family variant |

| | | | | |
|---|---|---|---|---|
| 3 | ETV5 | B | POLR2A | ETS family variant |
| 12 | ETV6 | B | POLR2A | ETS family variant |
| 6 | ETV7 | B | EZH2=POLR2A | ETS family variant |
| 7 | EVX1 | T | EZH2 | Homeobox |
| 2 | EVX2 | T | EZH2 | Homeobox |
| 17 | EZH1 | B | POLR2A | H3K27 methyl transferase |
| 7 | EZH2 | B | POLR2A | H3K27 methyl transferase |
| 14 | FOS | B | POLR2A | L-ZIP |
| 19 | FOSB | B | POLR2A | L-ZIP |
| 11 | FOSL1 | B | POLR2A + EZH2 | L-ZIP |
| 2 | FOSL2 | B | POLR2A | L-ZIP |
| 14 | FOXA1 | T | POLR2A > EZH2 | Forkhead |
| 20 | FOXA2 | T | EZH2 > POLR2A | Forkhead |
| 19 | FOXA3 | T | POLR2A | Forkhead |
| 15 | FOXB1 | T | EZH2 | Forkhead |
| 9 | FOXB2 | T | EZH2 | Forkhead |
| 6 | FOXC1 | B | POLR2A + EZH2 | Forkhead |
| 16 | FOXC2 | T | EZH2 > POLR2A | Forkhead |
| 1 | FOXD2 | T | EZH2 | Forkhead |
| 1 | FOXD3 | T | EZH2 > POLR2A | Forkhead |
| 9 | FOXD4 | T | EZH2 | Forkhead |
| 2 | FOXD4L1 | T | EZH2 | Forkhead-like |
| 9 | FOXD4L3 | T | neither | Forkhead-like |
| 9 | FOXD4L5 | T | neither | Forkhead-like |
| 9 | FOXD4L6 | T | EZH2 | Forkhead-like |
| 9 | FOXE1 | T | EZH2 | Forkhead |
| 1 | FOXE3 | T | EZH2 | Forkhead |
| 16 | FOXF1 | T | EZH2 | Forkhead |
| 6 | FOXF2 | T | EZH2 | Forkhead |
| 14 | FOXG1 | T | EZH2 | Forkhead |
| 8 | FOXH1 | T | EZH2 + POLR2A | Forkhead |
| 5 | FOXI1 | T | neither | Forkhead |
| 10 | FOXI2 | T | EZH2 | Forkhead |
| 17 | FOXJ1 | T | EZH2 > POLR2A | Forkhead |
| 12 | FOXJ2 | B | POLR2A | Forkhead |
| 1 | FOXJ3 | B | POLR2A | Forkhead |
| 17 | FOXK2 | B | POLR2A | Forkhead |
| 16 | FOXL1 | T | EZH2 | Forkhead |
| 3 | FOXL2 | T | EZH2 | Forkhead |
| 12 | FOXM1 | B | POLR2A | Forkhead |
| 17 | FOXN1 | T | POLR2A | Forkhead |
| 2 | FOXN2 | B | POLR2A- | Forkhead |
| 14 | FOXN3 | B | POLR2A | Forkhead |
| 12 | FOXN4 | T | EZH2 | Forkhead |
| 13 | FOXO1 | B | POLR2A | Forkhead |

| | | | | |
|---|---|---|---|---|
| 6 | FOXO3 | B | POLR2A | Forkhead |
| 23 | FOXO4 | B | POLR2A | Forkhead |
| 1 | FOXO6 | B | POLR2A + EZH2 | Forkhead |
| 3 | FOXP1 | B | POLR2A | Forkhead |
| 7 | FOXP2 | T | EZH2 + POLR2A | Forkhead |
| 23 | FOXP3 | B | POLR2A | Forkhead |
| 6 | FOXP4 | B | POLR2A | Forkhead |
| 6 | FOXQ1 | T | EZH2 + POLR2A | Forkhead |
| 11 | FOXR1 | T | neither | Forkhead |
| 23 | FOXR2 | T | neither | Forkhead |
| 20 | FOXS1 | T | POLR2A | Forkhead |
| 21 | GABPA | B | POLR2A | ETS family |
| 15 | GABPB1 | B | POLR2A | Similar to B2 |
| 1 | GABPB2 | B | POLR2A | Similar to B1 |
| 23 | GATA1 | T | POLR2A | ZNF |
| 3 | GATA2 | B | EZH2 > POLR2A | ZNF |
| 10 | GATA3 | T | EZH2 > POLR2A | ZNF |
| 8 | GATA4 | T | EZH2 | ZNF |
| 20 | GATA5 | T | EZH2 | ZNF |
| 18 | GATA6 | T | EZH2 | ZNF |
| 7 | GBX1 | T | EZH2 | Homeobox |
| 2 | GBX2 | T | EZH2 | Homeobox |
| 1 | GFI1 | T | EZH2 | ZNF |
| 9 | GFI1B | T | neither | ZNF |
| 12 | GLI1 | B | POLR2A | ZNF |
| 12 | GLI2 | B | POLR2A | ZNF |
| 12 | GLI3 | B | POLR2A | ZNF |
| 1 | GLIS1 | B | EZH2 | ZNF |
| 2 | GLIS2 | T | EZH2 | ZNF |
| 1 | GLIS3 | T | POLR2A + EZH2 | ZNF |
| 2 | GRHL1 | T | POLR2A | grainyhead |
| 8 | GRHL2 | T | POLR2A | grainyhead |
| 1 | GRHL3 | T | POLR2A + EZH2 | grainyhead |
| 14 | GSC | T | EZH2 | Homeobox bicoid |
| 22 | GSC2 | T | EZH2 | Homeobox |
| 13 | GSX1 | T | EZH2 | Homeobox-seq known |
| 4 | GSX2 | T | EZH2 | Homeobox |
| 5 | HAND1 | T | EZH2 | bHLH |
| 4 | HAND2 | T | EZH2 | bHLH |
| 3 | HES1 | B | POLR2A | bHLH |
| 1 | HES2 | T | EZH2 + POLR2A | bHLH |
| 1 | HES3 | T | EZH2 | bHLH |
| 1 | HES4 | B | EZH2 + POLR2A | bHLH |
| 1 | HES5 | T | EZH2 | bHLH |
| 17 | HES7 | T | EZH2 | bHLH |

| | | | | |
|---|---|---|---|---|
| 3 | HESX1 | B | POLR2A | Homeobox |
| 6 | HEY2 | B | EZH2 > POLR2A | bHLH |
| 1 | HEYL | B | EZH2 + POLR2A | bHLH |
| 17 | HIC1 | B | EZH2 + POLR2A | ZNF-BTB |
| 22 | HIC2 | B | POLR2A | ZNF-BTB |
| 14 | HIF1A | B | POLR2A | bHLH |
| 19 | HIF3A | B | POLR2A | bHLH |
| 6 | HIVEP1 | B | POLR2A | ZNF |
| 6 | HIVEP2 | B | POLR2A | ZNF |
| 1 | HIVEP3 | B | POLR2A | ZNF |
| 17 | HLF | B | POLR2A + EZH2 | bZIP |
| 1 | HLX | B | EZH2 | Homeobox |
| 4 | HMX1 | T | EZH2 | Homeobox |
| 10 | HMX2 | T | EZH2 | Homeobox |
| 10 | HMX3 | T | EZH2 | Homeobox |
| 12 | HNF1A | T | POLR2A | Homeobox-seq given |
| 17 | HNF1B | T | EZH2 + POLR2A | Homeobox- seq given |
| 8 | HNF4G | T | neither | NR subfamily |
| 7 | HOXA1 | T | EZH2 > POLR2A | Homeobox |
| 7 | HOXA10 | T | EZH2 > POLR2A | Homeobox |
| 7 | HOXA11 | T | EZH2=POLR2A | Homeobox |
| 7 | HOXA13 | T | EZH2 + POLR2A | Homeobox |
| 7 | HOXA2 | T | EZH2 > POLR2A | Homeobox |
| 7 | HOXA3 | T | EZH2 | Homeobox |
| 7 | HOXA4 | T | EZH2 | Homeobox |
| 7 | HOXA5 | T | EZH2 + POLR2A | Homeobox |
| 7 | HOXA6 | T | EZH2 + POLR2A | Homeobox |
| 7 | HOXA7 | T | EZH2 | Homeobox |
| 7 | HOXA9 | T | EZH2 | Homeobox |
| 17 | HOXB1 | T | EZH2 | Homeobox |
| 17 | HOXB13 | T | EZH2 | Homeobox |
| 17 | HOXB2 | B | EZH2 + POLR2A | Homeobox |
| 17 | HOXB3 | T | EZH2 > POLR2A | Homeobox |
| 17 | HOXB4 | B/T | EZH2 | Homeobox |
| 17 | HOXB5 | T | EZH2 > POLR2A | Homeobox |
| 17 | HOXB6 | T | EZH2 | Homeobox |
| 17 | HOXB7 | T | EZH2 > POLR2A | Homeobox |
| 17 | HOXB8 | T | EZH2 + POLR2A | Homeobox |
| 17 | HOXB9 | T | EZH2 + POLR2A | Homeobox |
| 12 | HOXC10 | T | EZH2 | Homeobox |
| 12 | HOXC11 | T | EZH2 | Homeobox |
| 12 | HOXC12 | T | EZH2 | Homeobox |
| 12 | HOXC13 | T | EZH2 | Homeobox |
| 12 | HOXC4 | T | EZH2 | Homeobox |
| 12 | HOXC5 | T | EZH2 | Homeobox |

| | | | | |
|---|---|---|---|---|
| 12 | HOXC6 | T | EZH2 | Homeobox |
| 12 | HOXC8 | T | EZH2 | Homeobox |
| 12 | HOXC9 | T | EZH2 | Homeobox |
| 2 | HOXD1 | T | EZH2 > POLR2A | Homeobox |
| 2 | HOXD10 | T | EZH2 | Homeobox |
| 2 | HOXD11 | T | EZH2 | Homeobox |
| 2 | HOXD12 | T | EZH2 | Homeobox |
| 2 | HOXD13 | T | EZH2 | Homeobox |
| 2 | HOXD3 | T | EZH2 | Homeobox |
| 2 | HOXD4 | T | EZH2 | Homeobox |
| 2 | HOXD8 | T | EZH2 | Homeobox |
| 2 | HOXD9 | T | EZH2 > POLR2A | Homeobox |
| 8 | HSF1 | B | POLR2A | Heat shock TF |
| 6 | HSF2 | B | POLR2A- | Heat shock TF |
| 16 | HSF4 | B | EZH2 + POLR2A | Heat shock TF |
| 17 | HSF5 | T | neither | Heat shock TF |
| 20 | ID1 | B | POLR2A | bHLH |
| 2 | ID2 | B | POLR2A | bHLH |
| 1 | ID3 | B | POLR2A | bHLH |
| 6 | ID4 | B | POLR2A | bHLH |
| 7 | IKZF1 | T | POLR2A > EZH2 | ZNF |
| 2 | IKZF2 | B | POLR2A | ZNF |
| 12 | IKZF4 | B | POLR2A | ZNF |
| 10 | IKZF5 | B | POLR2A | ZNF |
| 20 | INSM1 | T | EZH2 | ZNF |
| 14 | INSM2 | T | EZH2 | ZNF |
| 5 | IRF1 | B | POLR2A | HTH (W)5 repeat |
| 4 | IRF2 | B | POLR2A | HTH |
| 19 | IRF3 | B | POLR2A | HTH |
| 6 | IRF4 | T | POLR2A | HTH |
| 7 | IRF5 | B | POLR2A | HTH |
| 1 | IRF6 | T | POLR2A | HTH |
| 11 | IRF7 | B | POLR2A | HTH |
| 16 | IRF8 | B | POLR2A | HTH |
| 14 | IRF9 | B | POLR2A | HTH |
| 5 | IRX1 | T | EZH2 | Homeobox |
| 5 | IRX2 | T | POLR2A + EZH2 | Homeobox |
| 16 | IRX3 | T | EZH2 | Homeobox |
| 5 | IRX4 | T | EZH2 + POLR2A | Homeobox |
| 16 | IRX5 | T | EZH2 > POLR2A | Homeobox |
| 16 | IRX6 | T | EZH2 > POLR2A | Homeobox |
| 5 | ISL1 | T | EZH2 | LIM homeobox |
| 15 | ISL2 | T | EZH2 + POLR2A | LIM homeobox |
| 22 | ISX | T | POLR2A- | Homeobox |
| 1 | JUN | B | POLR2A | |

| | | | | |
|---|---|---|---|---|
| 19 | JUNB | B | POLR2A | |
| 19 | JUND | B | POLR2A | |
| 17 | KLF | B | POLR2A | ZNF |
| 19 | KLF1 | T | POLR2A | ZNF |
| 8 | KLF10 | B | POLR2A | ZNF |
| 2 | KLF11 | B | POLR2A | ZNF |
| 13 | KLF12 | B | POLR2A | ZNF |
| 15 | KLF13 | B | POLR2A | ZNF |
| 7 | KLF14 | T | EZH2 | ZNF |
| 3 | KLF15 | B | EZH2 + POLR2A | ZNF |
| 19 | KLF16 | B | POLR2A | ZNF |
| 1 | KLF17 | T | neither | ZNF |
| 19 | KLF2 | B | POLR2A + EZH2 | ZNF |
| 4 | KLF3 | B | POLR2A | ZNF |
| 13 | KLF5 | B | POLR2A | ZNF |
| 10 | KLF6 | B | POLR2A+ | ZNF |
| 2 | KLF7 | B | POLR2A | ZNF |
| 23 | KLF8 | B | POLR2A | ZNF |
| 9 | KLF9 | B | POLR2A | ZNF binds GC box |
| 10 | LBX1 | T | EZH2 | Homeobox |
| 2 | LBX2 | T | EZH2 + POLR2A | Homeobox |
| 10 | LCOR | B | neither | |
| 4 | LCORL | B | POLR2A | |
| 17 | LHX1 | T | EZH2 | LIM homeobox |
| 9 | LHX2 | T | EZH2 | LIM homeobox |
| 9 | LHX3 | T | EZH2 | LIM homeobox |
| 1 | LHX4 | B | EZH2 | LIM homeobox |
| 12 | LHX5 | T | EZH2 | LIM homeobox |
| 9 | LHX6 | T | EZH2 | LIM homeobox |
| 1 | LHX8 | T | EZH2 | LIM homeobox |
| 1 | LHX9 | T | EZH2 | LIM homeobox |
| 1 | LMX1A | T | EZH2 | LIM homeobox |
| 9 | LMX1B | T | EZH2 | LIM homeobox |
| 1 | MAEL | T | neither | |
| 16 | MAF | B | POLR2A + EZH2 | bZIP |
| 8 | MAFA | T | EZH2 | bZIP |
| 20 | MAFB | T | EZH2 | bZIP |
| 22 | MAFF | B | POLR2A | bZIP |
| 17 | MAFG | B | POLR2A | bZIP |
| 7 | MAFK | B | POLR2A | bZIP |
| 5 | MATR3 | B | POLR2A | Not TF? |
| 14 | MAX | B | POLR2A | bHLH |
| 16 | MAZ | B | POLR2A | ZNF |
| 15 | MEF2A | B | POLR2A | MADS box seq known |
| 19 | MEF2B | B | POLR2A | MADS box |

| | | | | |
|---|---|---|---|---|
| 5 | MEF2C | B | POLR2A | MADS box |
| 1 | MEF2D | B | POLR2A | MADS box |
| 2 | MEIS1 | B | EZH2 + POLR2A | Homeobox TALE |
| 15 | MEIS2 | B | POLR2A > EZH2 | Homeobox TALE |
| 19 | MEIS3 | T | EZH2 + POLR2A | Homeobox TALE |
| 1 | MIER1 | B | POLR2A | |
| 19 | MIER2 | B | POLR2A | |
| 5 | MIER3 | B | POLR2A | |
| 19 | MLLT1 | B | POLR2A | YEATS domain |
| 9 | MLLT3 | B | POLR2A + EZH2 | YEATS domain |
| 17 | MLX | B | POLR2A | bHLH-ZIP |
| 12 | MLXIP | B | POLR2A | bHLH-ZIP |
| 7 | MLXIPL | B | EZH2=POLR2A | bHLH-ZIP |
| 17 | MNT | B | POLR2A | bHLH-ZIP |
| 7 | MNX1 | T | EZH2 | Homeobox |
| 8 | MSC | B | EZH2 + POLR2A | bHLH |
| 4 | MSX1 | B | EZH2 + POLR2A | Homeobox |
| 5 | MSX2 | T | EZH2 | Homeobox |
| 2 | MXD1 | B | POLR2A | bHLH-ZIP |
| 5 | MXD3 | B | POLR2A | bHLH-ZIP |
| 4 | MXD4 | B | POLR2A | bHLH-ZIP |
| 10 | MXI1 | B | POLR2A | bHLH-ZIP |
| 6 | MYB | T | POLR2A + EZH2 | HTH seq known |
| 8 | MYBL1 | B | POLR2A | HTH |
| 20 | MYBL2 | T | POLR2A | HTH |
| 8 | MYC | B | POLR2A | bHLH |
| 1 | MYCL1 | T | POLR2A > EZH2 | bHLH |
| 2 | MYCN | T | EZH2 > POLR2A | bHLH |
| 12 | MYF5 | T | neither | bHLH |
| 12 | MYF6 | T | POLR2A | bHLH |
| 3 | MYNN | B | POLR2A | ZNF-BTB |
| 11 | MYOD1 | T | EZH2 | bHLH |
| 1 | MYOG | T | neither | bHLH |
| 20 | MYT1 | T | EZH2 | ZNF |
| 2 | MYT1L | T | neither | ZNF |
| 2 | NCOA1 | B | POLR2A | bHLH |
| 20 | NCOA3 | B | POLR2A | bHLH |
| 2 | NEUROD1 | T | EZH2 | bHLH |
| 12 | NEUROD4 | T | POLR2A | bHLH |
| 7 | NEUROD6 | T | EZH2 | bHLH |
| 5 | NEUROG1 | T | EZH2 | bHLH |
| 4 | NEUROG2 | T | EZH2 | bHLH |
| 10 | NEUROG3 | T | EZH2 | bHLH |
| 16 | NFAT5 | B | POLR2A | |
| 18 | NFATC1 | B | POLR2A + EZH2 | NFAT |

| | | | | |
|---|---|---|---|---|
| 16 | NFATC3 | B | POLR2A | |
| 14 | NFATC4 | B | POLR2A + EZH2 | |
| 12 | NFE2 | T | POLR2A | Seq known |
| 17 | NFE2L1 | B | POLR2A | |
| 2 | NFE2L2 | B | POLR2A | bZIP |
| 7 | NFE2L3 | B | POLR2A | bZIP |
| 1 | NFIA | B | POLR2A | |
| 9 | NFIB | B | POLR2A + EZH2 | |
| 19 | NFIC | B | POLR2A | NF-1 family |
| 9 | NFIL3 | B | POLR2A | |
| 19 | NFIX | B | EZH2>POLR2A | Seq known |
| 4 | NFKB1 | B | POLR2A | |
| 10 | NFKB2 | B | POLR2A | |
| 11 | NFRKB | B | POLR2A | |
| 9 | NFX1 | B | POLR2A | X box binding |
| 4 | NFXL1 | B | POLR2A | X box binding |
| 6 | NFYA | B | POLR2A- | Binds CCAAT |
| 12 | NFYB | B | POLR2A | Binds CCAAT |
| 1 | NFYC | B | POLR2A | Binds CCAAT |
| 1 | NHLH1 | T | POLR2A- | bHLH |
| 1 | NHLH2 | T | EZH2 | bHLH |
| 14 | NKX2-1 | T | EZH2 | Homeobox |
| 20 | NKX2-2 | T | EZH2 | Homeobox |
| 10 | NKX2-3 | T | EZH2 | Homeobox |
| 20 | NKX2-4 | T | EZH2 | Homeobox |
| 5 | NKX2-5 | T | EZH2 | Homeobox |
| 8 | NKX2-6 | T | EZH2 | Homeobox |
| 14 | NKX2-8 | T | EZH2 | Homeobox |
| 14 | NKX3-1 | T | EZH2 > POLR2A | Homeobox |
| 4 | NKX3-2 | T | EZH2 | Homeobox |
| 4 | NKX6-1 | T | EZH2 | Homeobox |
| 10 | NKX6-2 | T | EZH2 | Homeobox |
| 8 | NKX6-3 | T | EZH2 | Homeobox |
| 7 | NOBOX | T | neither | Homeobox |
| 2 | NOTO | T | EZH2 | Homeobox |
| 19 | NPAS1 | T | EZH2 | bHLH |
| 2 | NPAS2 | B | POLR2A | bHLH |
| 11 | NPAS4 | B | EZH2 + POLR2A | bHLH |
| 23 | NR0B1 | T | EZH2 | NR family |
| 17 | NR1D1 | B | POLR2A | NR family |
| 3 | NR1D2 | B | POLR2A | NR family |
| 19 | NR1H2 | B | POLR2A | NR family |
| 11 | NR1H3 | B | POLR2A | NR family |
| 12 | NR1H4 | T | POLR2A | NR family |
| 3 | NR1I2 | T | POLR2A | NR family |

| | | | | |
|---|---|---|---|---|
| 1 | NR1I3 | T | neither | NR family |
| 12 | NR2C1 | B | POLR2A | NR family |
| 3 | NR2C2 | B | POLR2A | NR family |
| 6 | NR2E1 | T | EZH2 | NR family |
| 15 | NR2E3 | T | neither | NR family |
| 5 | NR2F1 | B | EZH2 | NR family |
| 15 | NR2F2 | B | EZH2 | NR family |
| 19 | NR2F6 | B | POLR2A | NR family |
| 5 | NR3C1 | B | POLR2A | NR family |
| 4 | NR3C2 | B | POLR2A + EZH2 | NR family |
| 12 | NR4A1 | B | POLR2A | NR family |
| 2 | NR4A2 | B | POLR2A > EZH2 | NR family |
| 9 | NR4A3 | B/T | POLR2A + EZH2 | NR family |
| 9 | NR5A1 | T | EZH2 | NR family |
| 1 | NR5A2 | T | EZH2 + POLR2A | NR family |
| 9 | NR6A1 | B | POLR2A | NR family |
| 7 | NRF1 | B | POLR2A | |
| 14 | NRL | B | POLR2A + EZH2 | bZIP |
| 21 | OLIG2 | T | EZH2 | bHLH |
| 6 | OLIG3 | T | EZH2 | bHLH |
| 15 | ONECUT1 | T | EZH2 | Homeobox |
| 18 | ONECUT2 | T | EZH2 | Homeobox |
| 19 | ONECUT3 | T | EZH2 | Homeobox |
| 2 | OSR1 | T | EZH2 | |
| 8 | OSR2 | T | EZH2 | |
| 5 | OTP | T | EZH2 | Homeobox |
| 2 | OTX1 | T | EZH2 + POLR2A | Homeobox |
| 14 | OTX2 | T | EZH2 | Homeobox |
| 11 | OVOL1 | T | EZH2 | ZNF |
| 20 | OVOL2 | T | EZH2 | ZNF |
| 22 | PATZ1 | B | POLR2A | POZ-AT hook-ZNF |
| 12 | PAWR | B | POLR2A | bZIP |
| 20 | PAX1 | T | EZH2 | Paired box |
| 10 | PAX2 | T | EZH2 | Paired box |
| 2 | PAX3 | T | EZH2 | Paired box |
| 7 | PAX4 | T | EZH2 | Paired box |
| 9 | PAX5 | T | EZH2 | Paired box |
| 11 | PAX6 | T | EZH2 | Paired box |
| 1 | PAX7 | T | EZH2 | Paired box |
| 2 | PAX8 | T | POLR2A > EZH2 | Paired box |
| 14 | PAX9 | T | EZH2 | Paired box |
| 1 | PBX1 | B | POLR2A | Homeobox |
| 6 | PBX2 | B | POLR2A | Homeobox |
| 9 | PBX3 | B | POLR2A | Homeobox |
| 19 | PBX4 | B | POLR2A | Homeobox |

| | | | | |
|---|---|---|---|---|
| 12 | PHB2 | B | POLR2A | |
| 20 | PHF20 | B | POLR2A- | PHD finger ZNF? |
| 11 | PHOX2A | T | EZH2 > POLR2A | Homeobox |
| 4 | PHOX2B | T | EZH2 | Homeobox |
| 5 | PITX1 | T | EZH2 | Paired-like homeodomain |
| 4 | PITX2 | T | EZH2 | Paired-like homeodomain |
| 10 | PITX3 | T | EZH2 | Paired-like homeodomain |
| 21 | PKNOX1 | B | POLR2A | Homeobox |
| 11 | PKNOX2 | B | POLR2A + EZH2 | Homeobox |
| 8 | PLAG1 | B/T | EZH2 | ZNF |
| 6 | PLAGL1 | B | POLR2A + EZH2 | ZNF |
| 20 | PLAGL2 | B | POLR2A | ZNF |
| 3 | POU1F1 | T | neither | POU homeobox |
| 1 | POU2F1 | B | POLR2A | POU homeobox |
| 19 | POU2F2 | B | POLR2A | POU homeobox |
| 11 | POU2F3 | T | POLR2A > EZH2 | POU homeobox |
| 1 | POU3F1 | T | EZH2 | POU homeobox |
| 6 | POU3F2 | T | EZH2 | POU homeobox |
| 2 | POU3F3 | T | EZH2 | POU homeobox |
| 23 | POU3F4 | T | EZH2 | POU homeobox |
| 13 | POU4F1 | T | EZH2 | POU homeobox |
| 4 | POU4F2 | T | EZH2 | POU homeobox |
| 5 | POU4F3 | T | EZH2 | POU homeobox |
| 6 | POU5F1 | B | POLR2A | POU homeobox |
| 12 | POU6F1 | B | POLR2A | POU homeobox |
| 7 | POU6F2 | T | EZH2 | POU homeobox |
| 6 | PPARD | B | POLR2A | NR family |
| 3 | PPARG | B | POLR2A | NR family |
| 6 | PRDM1 | B | POLR2A | ZNF |
| 11 | PRDM10 | B | POLR2A | ZNF |
| 9 | PRDM12 | T | EZH2 | ZNF |
| 6 | PRDM13 | T | EZH2 | ZNF |
| 8 | PRDM14 | T | EZH2 | ZNF |
| 21 | PRDM15 | B | POLR2A | ZNF |
| 1 | PRDM16 | T | EZH2 | ZNF |
| 1 | PRDM2 | B | POLR2A | ZNF |
| 12 | PRDM4 | B | POLR2A | ZNF |
| 4 | PRDM5 | B | POLR2A | ZNF |
| 5 | PRDM6 | T | EZH2 | PR domain |
| 16 | PRDM7 | T | POLR2A | ZNF |
| 4 | PRDM8 | T/B | EZH2 | ZNF |
| 5 | PRDM9 | T | POLR2A- | ZNF |
| 2 | PREB | B | POLR2A | |
| 5 | PROP1 | T | EZH2 | Homeobox |
| 1 | PROX1 | T | EZH2 + POLR2A | Homeobox |

| | | | | |
|---|---|---|---|---|
| 14 | PROX2 | B | neither | Homeobox |
| 1 | PRRX1 | T | EZH2 | Homeobox |
| 9 | PRRX2 | T | EZH2 | Homeobox |
| 17 | RARA | B | POLR2A | NR family |
| 3 | RARB | B | POLR2A | NR family |
| 12 | RARG | B | POLR2A | NR family |
| 18 | RAX | T | EZH2 | Homeobox |
| 19 | RAX2 | T | neither | Homeobox |
| 7 | RBAK | B | POLR2A | ZNF |
| 4 | RBPJ | B | POLR2A | Binds NOTCH |
| 20 | RBPJL | T | EZH2 | Binds NOTCH |
| 9 | RC3H2 | B | POLR2A | ZNF |
| 14 | RCOR1 | B | POLR2A | Binds REST |
| 11 | RCOR2 | T | POLR2A + EZH2 | Binds REST |
| 1 | RCOR3 | B | POLR2A | Binds REST |
| 2 | REL | B | POLR2A | REL homology domain |
| 11 | RELA | B | POLR2A | REL homology domain |
| 19 | RELB | B | POLR2A | REL homology domain |
| 1 | RERE | B | POLR2A | RE repeats |
| 4 | REST | B | POLR2A | ZNF brain |
| 19 | RFX1 | B | POLR2A | RFX motif winged helix |
| 19 | RFX2 | B | POLR2A | RFX motif |
| 9 | RFX3 | B | POLR2A | RFX motif |
| 12 | RFX4 | T | EZH2 | RFX motif |
| 1 | RFX5 | B | POLR2A | RFX motif |
| 6 | RFX6 | T | EZH2 | RFX motif |
| 19 | RFXANK | B | POLR2A | |
| 23 | RHOXF1 | B | neither | Homeobox |
| 23 | RHOXF2 | T | neither | Homeobox |
| 15 | RORA | B | neither | NR1 family |
| 9 | RORB | T | EZH2 | NR1 family |
| 1 | RORC | T | POLR2A | NR1 family |
| 21 | RUNX1 | B | POLR2A | CBF subunit TGTGGT |
| 6 | RUNX2 | B | POLR2A | CBF subunit |
| 1 | RUNX3 | B | POLR2A | CBF subunit |
| 6 | RXRB | B | POLR2A | NR family |
| 1 | RXRG | T | EZH2 | NR family |
| 16 | SALL1 | T | EZH2 | ZNF |
| 14 | SALL2 | T | POLR2A | ZNF |
| 18 | SALL3 | T | EZH2 | ZNF |
| 20 | SALL4 | T | POLR2A | ZNF |
| 3 | SATB1 | B | EZH2 + POLR2A | Homeobox |
| 2 | SATB2 | T | EZH2 + POLR2A | Homeobox |
| 8 | SCRT1 | T | EZH2 | ZNF |
| 20 | SCRT2 | T | EZH2 | ZNF |

| | | | | |
|---|---|---|---|---|
| 6 | SIM1 | T | EZH2 | bHLH |
| 21 | SIM2 | T | EZH2 | bHLH |
| 14 | SIX1 | T | EZH2 | Homeobox |
| 2 | SIX2 | T | EZH2 + POLR2A | Homeobox |
| 2 | SIX3 | T | EZH2 | Homeobox |
| 14 | SIX4 | B | POLR2A | Homeobox |
| 19 | SIX5 | B | POLR2A | Homeobox |
| 14 | SIX6 | T | EZH2 | Homeobox |
| 1 | SKI | B | POLR2A | |
| 3 | SKIL | B | POLR2A | |
| 4 | SMAD1 | B | POLR2A | SMAD family |
| 18 | SMAD2 | B | POLR2A | SMAD family |
| 15 | SMAD3 | B | POLR2A | SMAD family |
| 18 | SMAD4 | B | POLR2A | SMAD family |
| 5 | SMAD5 | B | POLR2A | SMAD family |
| 13 | SMAD9 | B | EZH2 + POLR2A | SMAD family |
| 20 | SNAI1 | B | POLR2A | ZNF |
| 8 | SNAI2 | B | POLR2A>EZH2 | ZNF |
| 16 | SNAI3 | B | POLR2A | ZNF |
| 9 | SNAPC4 | B | POLR2A | SNAP family |
| 13 | SOX1 | T | EZH2 | SOX family |
| 22 | SOX10 | T | POLR2A + EZH2 | SOX family |
| 2 | SOX11 | T | EZH2 | SOX family |
| 20 | SOX12 | B | POLR2A | SOX family |
| 1 | SOX13 | B | POLR2A | SOX family |
| 3 | SOX14 | T | EZH2 | SOX family |
| 17 | SOX15 | B | POLR2A | SOX family |
| 8 | SOX17 | T | EZH2 > POLR2A | SOX family |
| 20 | SOX18 | B | EZH2 + POLR2A | SOX family |
| 3 | SOX2 | T | EZH2 + POLR2A | SOX family |
| 13 | SOX21 | T | EZH2 | SOX family |
| 23 | SOX3 | T | EZH2 + POLR2A | SOX family |
| 5 | SOX30 | T | POLR2A- | SOX family |
| 6 | SOX4 | B | POLR2A > EZH2 | SOX family |
| 12 | SOX5 | B | POLR2A | SOX family |
| 11 | SOX6 | B | POLR2A | SOX family |
| 8 | SOX7 | B | EZH2 + POLR2A | SOX family |
| 16 | SOX8 | T | EZH2 | SOX family |
| 17 | SOX9 | B | POLR2A > EZH2 | SOX family |
| 12 | SP1 | B | POLR2A | ZNF |
| 2 | SP100 | B | POLR2A | ? not TF? |
| 2 | SP110 | B | POLR2A | Nuclear body protein |
| 17 | SP2 | B | POLR2A | Sp family |
| 2 | SP3 | B | POLR2A | Sp family |
| 7 | SP4 | B | POLR2A | Sp family |

| | | | | |
|---|---|---|---|---|
| 2 | SP5 | T | POLR2A + EZH2 | Sp family |
| 17 | SP6 | T | EZH2 | Sp family |
| 12 | SP7 | T | EZH2 | Sp family |
| 7 | SP8 | T | EZH2 | Sp family |
| 11 | SPI1 | B | POLR2A | ETS family |
| 19 | SPIB | T | POLR2A + EZH2 | ETS family |
| 12 | SPIC | T | neither | ETS family |
| 17 | SREBF1 | B | POLR2A | bHLH |
| 22 | SREBF2 | B | POLR2A | bHLH |
| 6 | SRF | B | POLR2A | MADS box |
| 24 | SRY | T | neither | HMG box |
| 2 | STAT1 | B | POLR2A | STAT family |
| 12 | STAT2 | B | POLR2A | STAT family |
| 17 | STAT3 | B | POLR2A | STAT family |
| 2 | STAT4 | B | POLR2A- | STAT family |
| 17 | STAT5A | B | POLR2A | STAT family |
| 17 | STAT5B | B | POLR2A | STAT family |
| 12 | STAT6 | B | POLR2A | STAT family |
| 1 | TAL1 | B | EZH2 | bHLH |
| 9 | TAL2 | T | neither | bHLH |
| 11 | TBX10 | T | POLR2A | T box TCACACCT |
| 1 | TBX15 | T | EZH2 | T box early embryo |
| 6 | TBX18 | T | EZH2 | T box |
| 1 | TBX19 | B | POLR2A- | T box |
| 17 | TBX2 | B | EZH2 >> POLR2A | T box |
| 7 | TBX20 | T | EZH2 | T box |
| 17 | TBX21 | T | EZH2+POLR2A | T box |
| 23 | TBX22 | T | EZH2 | T box |
| 12 | TBX3 | B | EZH2 > POLR2A | T box |
| 17 | TBX4 | T | EZH2 | T box |
| 12 | TBX5 | T | EZH2 | T box |
| 16 | TBX6 | B | POLR2A | T box |
| 6 | TBXT | T | EZH2 | T box |
| 15 | TCF12 | B | POLR2A | bHLH |
| 20 | TCF15 | T | EZH2 > POLR2A | bHLH |
| 6 | TCF19 | B | POLR2A | ZNF |
| 22 | TCF20 | B | neither | NR? |
| 6 | TCF21 | T | EZH2 | bHLH |
| 2 | TCF23 | T | POLR2A | bHLH |
| 8 | TCF24 | T | EZH2 | bHLH |
| 16 | TCF25 | B | POLR2A | bHLH |
| 19 | TCF3 | B | POLR2A | bHLH |
| 18 | TCF4 | B | POLR2A | bHLH |
| 5 | TCF7 | B | POLR2A | HMG box |
| 2 | TCF7L1 | B | POL + EZH2 | HMG box |

| | | | | |
|---|---|---|---|---|
| 10 | TCF7L2 | B | POLR2A | HMG box |
| 20 | TCFL5 | B | POLR2A | bHLH |
| 11 | TEAD1 | B | POLR2A | TEA domain CATTCC[AT] |
| 19 | TEAD2 | B | POLR2A | TEA domain |
| 6 | TEAD3 | B | POLR2A | TEA domain |
| 12 | TEAD4 | B | POLR2A | TEA domain |
| 6 | TFAP2A | T | EZH2 + POLR2A | AP-2 group |
| 6 | TFAP2B | T | EZH2 | AP-2 group |
| 20 | TFAP2C | T | EZH2>POLR2A | AP-2 group |
| 6 | TFAP2D | T | EZH2 | AP-2 group |
| 1 | TFAP2E | B | POLR2A + EZH2 | AP-2 group |
| 16 | TFAP4 | B | POLR2A | bHLH |
| 12 | TFCP2 | B | POLR2A | |
| 23 | TFCP2L1 | B/T | POLR2A/EZH2 | |
| 13 | TFDP1 | B | POLR2A | E2F dimerization |
| 3 | TFDP2 | B | POLR2A | E2F dimerization |
| 23 | TFDP3 | T | neither | E2F dimerization |
| 23 | TFE3 | B | POLR2A | bHLH |
| 6 | TFEB | B | POLR2A | bHLH |
| 7 | TFEC | T | neither | bHLH |
| 18 | TGIF1 | B | POLR2A | Homeobox (TALE) |
| 23 | TGIF2LX | T | POLR2A- | Homeobox |
| 24 | TGIF2LY | T | neither | Homeobox |
| 15 | THAP10 | B | POLR2A | THAP domain ZNF |
| 1 | THAP3 | B | POLR2A | THAP domain ZNF |
| 17 | THRA | B | POLR2A | NR family |
| 3 | THRB | B | POLR2A + EZH2 | NR family |
| 10 | TLX1 | T | EZH2 | Homeobox (NK-like) |
| 2 | TLX2 | T | EZH2 | Homeobox |
| 5 | TLX3 | T | EZH2 | Homeobox |
| 8 | TOX | T | POLR2A + EZH2 | HMG box |
| 20 | TOX2 | B | EZH2 + POLR2A | HMG box |
| 16 | TOX3 | T | POLR2A + EZH2 | HMG box |
| 14 | TOX4 | B | POLR2A | HMG box |
| 17 | TP53 | B | POLR2A | |
| 3 | TP63 | T | POLR2A | |
| 1 | TP73 | T | POLR2A | |
| 13 | TSC22D1 | B | POLR2A | L-ZIP TSC22 domain |
| 23 | TSC22D3 | B | POLR2A | L-ZIP TSC22 domain |
| 7 | TSC22D4 | B | POLR2A | L-ZIP TSC22 domain |
| 18 | TSHZ1 | B | POLR2A | ZNF homeobox |
| 20 | TSHZ2 | T | POLR2A | ZNF homeobox |
| 19 | TSHZ3 | T | POLR2A = EZH2 | ZNF homeobox |
| 3 | UBP1 | B | POLR2A | |
| 17 | UBTF | B | POLR2A | HMG box |

| | | | | |
|---|---|---|---|---|
| 7 | UNCX | T | EZH2 | Homeobox |
| 16 | UNKL | B | POLR2A | ZNF |
| 1 | USF1 | B | POLR2A | |
| 19 | USF2 | B | POLR2A | |
| 10 | VAX1 | T | EZH2 | Homeobox |
| 2 | VAX2 | T/B | EZH2 > POLR2A | Homeobox |
| 20 | VSX1 | T | EZH2 | Homeobox |
| 14 | VSX2 | T | EZH2 | Homeobox |
| 1 | YBX1 | B | POLR2A | Y-box |
| 1 | YBX2 | B | POLR2A | Y-box |
| 12 | YBX3 | B | POLR2A | Y-box |
| 14 | YY1 | B | POLR2A | ZNF |
| 23 | YY2 | B | neither | ZNF |
| 23 | ZBED1 | B | POLR2A | ZNF |
| 22 | ZBED4 | B | POLR2A > EZH2 | ZNF |
| 14 | ZBTB1 | B | POLR2A- | ZNF-BTB |
| 3 | ZBTB11 | B | POLR2A | ZNF-BTB |
| 18 | ZBTB14 | B | POLR2A | ZNF-BTB |
| 11 | ZBTB16 | B | EZH2 > POLR2A | ZNF-BTB |
| 1 | ZBTB17 | B | POLR2A | ZNF-BTB |
| 1 | ZBTB18 | B | POLR2A | ZNF |
| 6 | ZBTB2 | B | POLR2A | ZNF-BTB |
| 3 | ZBTB20 | B | POLR2A | ZNF-BTB |
| 21 | ZBTB21 | B | POLR2A | ZNF |
| 6 | ZBTB24 | B | POLR2A | ZNF-BTB |
| 19 | ZBTB32 | T | POLR2A | ZNF-BTB |
| 23 | ZBTB33 | B | POLR2A | ZNF-BTB |
| 9 | ZBTB34 | B | POLR2A | ZNF-BTB |
| 3 | ZBTB38 | B | POLR2A- | ZNF-BTB |
| 12 | ZBTB39 | B | POLR2A | ZNF-BTB |
| 17 | ZBTB4 | B | POLR2A | ZNF-BTB |
| 1 | ZBTB41 | B | POLR2A | ZNF-BTB |
| 9 | ZBTB43 | B | POLR2A | ZNF-BTB |
| 19 | ZBTB45 | B | POLR2A | ZNF-BTB |
| 20 | ZBTB46 | B | EZH2 + POLR2A | ZNF-BTB |
| 3 | ZBTB47 | B | POLR2A | ZNF-BTB |
| 1 | ZBTB48 | B | POLR2A | ZNF-BTB |
| 9 | ZBTB5 | B | POLR2A | ZNF-BTB |
| 19 | ZBTB7A | B | POLR2A | ZNF-BTB |
| 1 | ZBTB7B | B | POLR2A | ZNF-BTB |
| 1 | ZC3H11A | B | POLR2A | ZNF E-box homeobox |
| 2 | ZC3H15 | B | POLR2A | ZNF E-box homeobox |
| 8 | ZC3H3 | B | POLR2A | ZNF E-box homeobox |
| 10 | ZEB1 | B | POLR2A | ZNF + homeobox |
| 2 | ZEB2 | B | EZH2 + POLR2A | ZNF + homeobox |

| | | | | |
|---|---|---|---|---|
| 8 | ZFAT | B | POLR2A | ZNF AT-Hook domain |
| 14 | ZFHX2 | B | POLR2A | ZNF homeobox |
| 14 | ZFHX2 | B | POLR2A | ZNF |
| 16 | ZFHX3 | B | EZH2 + POLR2A | ZNF homeobox |
| 9 | ZFP37 | B | EZH2 > POLR2A | ZNF |
| 20 | ZFP64 | B | POLR2A | ZNF |
| 16 | ZFPM1 | B | POLR2A | ZNF FOG |
| 8 | ZFPM2 | B | POLR2A > EZH2 | ZNF FOG |
| 23 | ZFX | B | POLR2A | ZNF X-linked |
| 24 | ZFY | B | POLR2A | ZNF Y linked |
| 14 | ZFYVE26 | B | POLR2A | ZNF FYVE type |
| 8 | ZHX1 | B | POLR2A | ZNF + homeobox |
| 8 | ZHX2 | B | EZH2 + POLR2A | ZNF + homeobox |
| 20 | ZHX3 | B | POLR2A | |
| 3 | ZIC1 | T | EZH2 | ZNF |
| 13 | ZIC2 | T | POLR2A+ EZH2+ | ZNF |
| 23 | ZIC3 | T | EZH2 | ZNF |
| 3 | ZIC4 | T | EZH2 | ZNF |
| 13 | ZIC5 | T | EZH2 | ZNF |
| 19 | ZKSCAN1 | T | POLR2A | ZNF + KRAB +SCAN |
| 16 | ZKSCAN2 | B | POLR2A | ZNF + KRAB +SCAN |
| 6 | ZKSCAN3 | B | POLR2A | ZNF + KRAB +SCAN |
| 11 | ZKSCAN4 | B | POLR2A | ZNF + KRAB +SCAN |
| 7 | ZKSCAN5 | B | POLR2A | ZNF + KRAB +SCAN |
| 23 | ZMAT1 | B | POLR2A | ZNF Martin type |
| 3 | ZMAT3 | B | POLR2A | ZNF Martin type |
| 12 | ZNF10 | B | POLR2A | ZNF |
| 7 | ZNF12 | B | POLR2A | ZNF |
| 5 | ZNF131 | B | POLR2A | ZNF |
| 19 | ZNF134 | B | POLR2A | ZNF |
| 19 | ZNF136 | B | POLR2A | ZNF |
| 11 | ZNF143 | B | POLR2A | ZNF |
| 19 | ZNF146 | B | POLR2A | ZNF |
| 3 | ZNF148 | B | POLR2A | ZNF |
| 23 | ZNF157 | T | neither | ZNF |
| 8 | ZNF16 | B | POLR2A | ZNF |
| 16 | ZNF174 | B | POLR2A | ZNF |
| 19 | ZNF175 | B | POLR2A | ZNF |
| 6 | ZNF184 | B | POLR2A | ZNF |
| 11 | ZNF202 | B | POLR2A | ZNF |
| 20 | ZNF217 | B | POLR2A | ZNF |
| 14 | ZNF219 | B | POLR2A | ZNF |
| 19 | ZNF224 | B | POLR2A | ZNF |
| 18 | ZNF24 | B | POLR2A | ZNF |
| 8 | ZNF251 | B | POLR2A | ZNF |

| | | | | |
|---|---|---|---|---|
| 19 | ZNF256 | B | POLR2A | ZNF |
| 19 | ZNF260 | B | POLR2A | ZNF |
| 16 | ZNF276 | B | POLR2A | ZNF |
| 22 | ZNF280A | T | neither | ZNF |
| 1 | ZNF281 | B | POLR2A | ZNF |
| 7 | ZNF282 | B | POLR2A | ZNF |
| 6 | ZNF292 | B | POLR2A | ZNF |
| 19 | ZNF296 | B | POLR2A | ZNF |
| 5 | ZNF300 | B | POLR2A | ZNF |
| 19 | ZNF304 | B | POLR2A | ZNF |
| 16 | ZNF319 | B | POLR2A | ZNF |
| 10 | ZNF32 | B | POLR2A | ZNF |
| 6 | ZNF322A | B | POLR2A | ZNF |
| 19 | ZNF333 | B | POLR2A | ZNF |
| 20 | ZNF335 | B | POLR2A | ZNF |
| 20 | ZNF341 | B | POLR2A | ZNF |
| 19 | ZNF350 | B | POLR2A | ZNF |
| 5 | ZNF354A | B | POLR2A | ZNF |
| 5 | ZNF354B | B | POLR2A | ZNF |
| 5 | ZNF354C | B | POLR2A | ZNF |
| 1 | ZNF362 | B | POLR2A | ZNF |
| 10 | ZNF365 | T | EZH2 | ZNF |
| 5 | ZNF366 | B | POLR2A | ZNF |
| 9 | ZNF367 | B | POLR2A | ZNF |
| 10 | ZNF37A | B | POLR2A | ZNF |
| 19 | ZNF382 | B | POLR2A | ZNF |
| 12 | ZNF384 | B | POLR2A | ZNF |
| 7 | ZNF394 | B | POLR2A | ZNF |
| 7 | ZNF398 | B | POLR2A | ZNF |
| 14 | ZNF410 | B | POLR2A | ZNF |
| 19 | ZNF415 | B | POLR2A | ZNF |
| 19 | ZNF418 | B | POLR2A | ZNF |
| 16 | ZNF423 | B | POLR2A | ZNF |
| 7 | ZNF425 | B | POLR2A | ZNF |
| 19 | ZNF426 | B | POLR2A | ZNF |
| 19 | ZNF430 | B | POLR2A | ZNF |
| 19 | ZNF431 | B | POLR2A | ZNF |
| 10 | ZNF438 | B | POLR2A | ZNF |
| 19 | ZNF444 | B | POLR2A | ZNF |
| 3 | ZNF445 | B | POLR2A | ZNF |
| 7 | ZNF467 | B | POLR2A | ZNF |
| 19 | ZNF480 | B | POLR2A | ZNF |
| 10 | ZNF488 | T | POLR2A | ZNF |
| 1 | ZNF496 | B | POLR2A | ZNF |
| 4 | ZNF509 | B | POLR2A | ZNF |

| | | | | |
|---|---|---|---|---|
| 9 | ZNF510 | B | POLR2A | ZNF |
| 2 | ZNF513 | B | POLR2A | ZNF |
| 18 | ZNF516 | B | POLR2A | ZNF |
| 18 | ZNF521 | B | EZH2 | ZNF |
| 19 | ZNF536 | T | POLR2A | ZNF |
| 19 | ZNF551 | B | POLR2A | ZNF |
| 19 | ZNF568 | B | POLR2A | ZNF |
| 1 | ZNF593 | B | POLR2A | ZNF |
| 16 | ZNF598 | B | POLR2A | ZNF |
| 19 | ZNF606 | B | POLR2A | ZNF |
| 5 | ZNF608 | B | POLR2A | ZNF |
| 15 | ZNF609 | B | POLR2A | ZNF |
| 5 | ZNF622 | B | POLR2A | ZNF |
| 19 | ZNF628 | B | POLR2A | ZNF |
| 2 | ZNF638 | B | POLR2A | ZNF |
| 12 | ZNF641 | B | POLR2A | ZNF |
| 1 | ZNF642 | B | POLR2A | ZNF |
| 1 | ZNF644 | B | POLR2A | ZNF |
| 19 | ZNF649 | B | POLR2A | ZNF |
| 17 | ZNF652 | B | POLR2A | ZNF |
| 19 | ZNF653 | B | POLR2A | ZNF |
| 9 | ZNF658 | B | POLR2A | ZNF |
| 1 | ZNF683 | T | neither | ZNF |
| 1 | ZNF691 | B | POLR2A | ZNF |
| 19 | ZNF699 | B | POLR2A | ZNF |
| 8 | ZNF703 | B | POLR2A | ZNF |
| 8 | ZNF704 | B | POLR2A | ZNF |
| 8 | ZNF706 | B | POLR2A | ZNF |
| 15 | ZNF710 | B | POLR2A | ZNF |
| 23 | ZNF711 | B | POLR2A | ZNF |
| 12 | ZNF740 | B | POLR2A | ZNF |
| 7 | ZNF746 | B | POLR2A | ZNF |
| 17 | ZNF750 | T | POLR2A | ZNF |
| 7 | ZNF777 | B | POLR2A | ZNF |
| 9 | ZNF782 | B | POLR2A | ZNF |
| 7 | ZNF786 | B | POLR2A | ZNF |
| 19 | ZNF8 | B | POLR2A | ZNF |
| 16 | ZNF821 | B | POLR2A | ZNF |
| 19 | ZNF829 | B | POLR2A | ZNF |
| 17 | ZNF830 | B | POLR2A | ZNF |
| 20 | ZNF831 | T | POLR2A | ZNF |
| 19 | ZNF837 | B | POLR2A | ZNF |
| 19 | ZNF845 | B | POLR2A | ZNF |
| 19 | ZNF91 | B | POLR2A | ZNF |
| 19 | ZNF93 | B | POLR2A | ZNF |

| | | | | |
|---|---|---|---|---|
| 19 | ZSCAN1 | T | POLR2A | ZNF + SCAN |
| 16 | ZSCAN10 | T | POLR2A | ZNF + SCAN |
| 6 | ZSCAN12 | B | neither | ZNF + SCAN |
| 6 | ZSCAN16 | B | POLR2A | ZNF + SCAN |
| 15 | ZSCAN2 | B | POLR2A- | ZNF + SCAN |
| 19 | ZSCAN20 | B | POLR2A | ZNF + SCAN |
| 7 | ZSCAN21 | B | POLR2A | ZNF + SCAN |
| 19 | ZSCAN22 | B | POLR2A | ZNF + SCAN |
| 6 | ZSCAN23 | B | POLR2A | ZNF + SCAN |
| 15 | ZSCAN29 | B | POLR2A | ZNF + SCAN |
| 19 | ZSCAN4 | T | neither | ZNF + SCAN |
| 19 | ZSCAN5A | T | POLR2A | ZNF + SCAN |
| 23 | ZXDA | B | POLR2A | ZNF X-linked 10X ZNF |
| 23 | ZXDB | B | POLR2A | ZNF X-linked |
